# Dual functionality of the TasA amyloid protein in *Bacillus* physiology and fitness on the phylloplane

**DOI:** 10.1101/651356

**Authors:** Jesús Cámara-Almirón, Yurena Navarro, M. Concepción Magno-Pérez-Bryan, Carlos Molina-Santiago, John R. Pearson, Luis Díaz-Martínez, Antonio de Vicente, Alejandro Pérez-García, Diego Romero

**Author notes:** These authors contributed equally to this work (Jesús Cámara-Almirón, Yurena Navarro).

## Abstract

Bacteria can form biofilms that consist of multicellular communities embedded in an extracellular matrix (ECM). Previous studies have demonstrated that genetic pathways involved in biofilm formation are activated under a variety of environmental conditions to enhance bacterial fitness; however, the functions of the individual ECM components are still poorly understood. In *Bacillus subtilis*, the main protein component of the ECM is the functional amyloid TasA. In this study, we demonstrate that beyond their well-known defect in biofilm formation, *ΔtasA* cells also exhibit a range of cytological symptoms indicative of excessive cellular stress, including DNA damage accumulation, changes in membrane potential, higher susceptibility to oxidative stress, and alterations in membrane dynamics. Collectively, these events can lead to increased programmed cell death in the colony. We show that these major physiological changes in *ΔtasA* cells are likely independent of the structural role of TasA during amyloid fiber formation in the ECM. The presence of TasA in cellular membranes, which would place it in proximity to functional membrane microdomains, and mislocalization of the flotillin-like protein FloT in *ΔtasA* cells, led us to propose a role for TasA in the stabilization of membrane dynamics as cells enter stationary phase. We found that these alterations caused by the absence of TasA impair the survival, colonization and competition of *Bacillus* cells on the phylloplane. Taken together, our results allow the separation of two complementary roles of this functional amyloid protein: i) structural functions during ECM assembly and interactions with plants, and ii) a physiological function in which TasA, via its localization to the cell membrane, stabilizes membrane dynamics and supports more effective cellular adaptation to environmental cues.

## Introduction

In response to a wide range of environmental factors^1, 2^, some bacterial species establish complex communities called biofilms^3^. To do so, planktonic cells initiate a transition into a sedentary lifestyle and trigger a cell differentiation program that leads to two distinctive features: (1) a division of labor, in which different subpopulations of cells are dedicated to covering different processes needed to maintain the viability of the community^4, 5^, and (2) the secretion of a battery of molecules that assemble the extracellular matrix (ECM)^3, 6^. Like in eukaryotic tissues, the bacterial ECM is a dynamic structure that supports cellular adhesion and regulates the flux of signals to ensure cell differentiation, both of which are key ECM functions in biofilms^7, 8^. The tissue-like structure of biofilms also provides stability and serves as an interface with the external environment, working as a formidable physicochemical barrier against external assaults^9–11^. In eukaryotic cells, the ECM plays an important role in signaling^12, 13^ and has been described as a reservoir for the localization and concentration of growth factors, which in turn form gradients that are critical for the establishment of developmental patterning during morphogenesis^14, 15, 16^. Interestingly, in senescent cells, partial loss of the ECM as well as rearrangement of its components via an interplay between the activities of various matrix metalloproteases (MMPs) and tissue-specific MMP inhibitors can influence cell fate, e.g., by activating the apoptotic program^17, 18^. In both eukaryotes and prokaryotes, senescence involves global changes in cellular physiology, and in some microbes, this process begins with the entry of the cells into stationary phase^19–21^. At this stage, the rate of cell division slows^21^, the molecular machinery adapts to increase cellular resistance and the respiration and primary metabolism shift to fermentative pathways and to the production of secondary metabolites, respectively^22^. This process triggers a response typified by molecular mechanisms evolved to overcome environmental adversities and to ensure survival, including the activation of general stress response genes^23, 24^, a shift to anaerobic respiration^22^, enhanced DNA repair^25^, and induction of pathways for the metabolism of alternative nutrient sources or sub-products of primary metabolism^26^.

Studies of *Bacillus subtilis* biofilms have contributed to our understanding of the intricate developmental program that underlies biofilm formation^27–30^. External receptors oversee the sensing of a myriad of signals that must be properly integrated via an interconnected network of genetic cascades that end with the expression and secretion of ECM components accompanied by other cellular changes. Currently, the *B. subtilis* ECM is known to consist mainly of exopolysaccharide (EPS) and the TasA and BslA proteins^27^. Mutations affecting any of these components lead to different morphological phenotypes, reflecting their complementary functions in establishing the final architecture of the biofilm. The EPS acts as the adhesive element of the biofilm cells at the cell-to-surface interface, which is important for biofilm attachment^31^, and BslA is a hydrophobin that forms a thin external hydrophobic layer and is the main factor that confers hydrophobic properties to biofilms^32^. Both structural factors contribute to maintain the defense function performed by the ECM^11, 32^. TasA is a functional amyloid protein that forms resistant fibers that confer structural stability to biofilms^33, 34^. Additional proteins are needed for the polymerization of these fibers: TapA appears to favor the transition of TasA into the fiber state, and the signal peptidase SipW processes both proteins into their mature forms^35, 36^. Amyloids are widespread in nature, and studies in various experimental systems are expanding our view of the functions of this heterogeneous family of proteins. The ability of amyloids to transition from monomers into fibers represents structural, biochemical and functional versatility that microbes exploit in different contexts and for different purposes, e.g., the formation of adhesins and other ECM components, virulence expression, and competition with other organisms^37^.

Previous studies have demonstrated that the genetic pathways involved in biofilm formation are active during the interaction of several microbial species with plants^38, 39^. In *B. subtilis,* the lipopeptide surfactin acts as a self-trigger of biofilm formation on the melon phylloplane, which is connected with the suppressive activity of *B. subtilis* against phytopathogenic fungi^40^. These findings led us to hypothesize that the ECM makes a major contribution to the ecology of *B. subtilis* in the poorly explored phyllosphere. Our study of the ecology of *B. subtilis* NCIB3610-derived strains carrying single mutations in different ECM components in the phyllosphere highlights the role of TasA in bacteria-plant interactions. Moreover, the increased production of secondary metabolites by a *tasA* mutant strain on plant leaves revealed a complementary role for TasA in the stabilization of the bacteria’s physiology. In *ΔtasA* cells, gene expression changes and dynamic cytological alterations affect membrane potential, adaptation to oxidative stress and membrane functionality and dynamics, which eventually lead to a premature increase in programmed cell death (PCD) within the colony. In addition, two complementary pieces of evidence prove that these alterations are independent of the structural role of TasA in ECM assembly: i) we report that TasA is associated with the detergent-resistant fraction of the cell membrane (DRM) and that its absence leads to changes in membrane dynamics, as indicated by the mislocalization of the flotillin-like protein FloT, which is involved in the regulation of many of these physiological functions; and ii) ectopic expression of a mutated TasA protein in a *tasA* null mutant background fails to restore the strain’s ability to form biofilms and antagonize a phytopathogenic fungus on plants, while it does restore the mutant strain’s ability to maintain the physiological status of the cells. All these results indicate that these two complementary roles of TasA, both as part of the ECM and in regulating cell membrane dynamics, are important to preserve cell viability within the colony and for the ecological fitness of *B. subtilis* in the phylloplane.

## Results

### TasA contributes to the fitness of *Bacillus* on the phylloplane

Surfactin, a member of a subfamily of lipopeptides produced by *B. subtilis* and related species, contributes to the multicellularity of biofilms by triggering a potassium leakage that is detected by the sensor kinase KinC, which ultimately activates the expression of the *eps* and *tapA* operons required for biofilm formation^41^. Reflecting the contribution made by surfactin to multicellularity, we previously reported how a mutant strain defective for lipopeptide production showed impaired biofilm assembly on the phylloplane^40^. These two observations led us to evaluate the specific contributions made by the ECM structural components TasA and the EPS to *B. subtilis* fitness on melon leaves. Although not directly linked to the surfactin-activated regulatory pathway, we also studied the gene encoding the hydrophobin protein BslA (another important ECM component). A *tasA* mutant strain (*ΔtasA*) is defective in the initial cell attachment to plant surfaces (4 hours and two days post-inoculation) (Fig. 1A, top and Fig. S1A). As expected, based on their structural functions, all of the matrix mutants showed reduced survival; however, the population of *ΔtasA* cells continuously and steadily decreased over time compared to the populations of *eps* or *bslA* mutant cells (Fig. 1B and Fig. S1B). Examination of plants inoculated with the wild type strain (WT) or with the *ΔtasA* strain via scanning electron microscopy (SEM) revealed variability in the colonization patterns of the strains. WT cells assembled in ordered and compact colonies, with the cells embedded in a network of extracellular material (Fig. 1C, top). In contrast, the *ΔtasA* cells were prone to irregular distribution as large masses of cells on the leaves (Fig. 1C, bottom). Finally, *eps* and *bslA* mutant cells formed flat colonies (Fig. S2A) with the same colonization defects observed in the *tasA* mutant cells (Fig. S1C).

**Figure 1.**
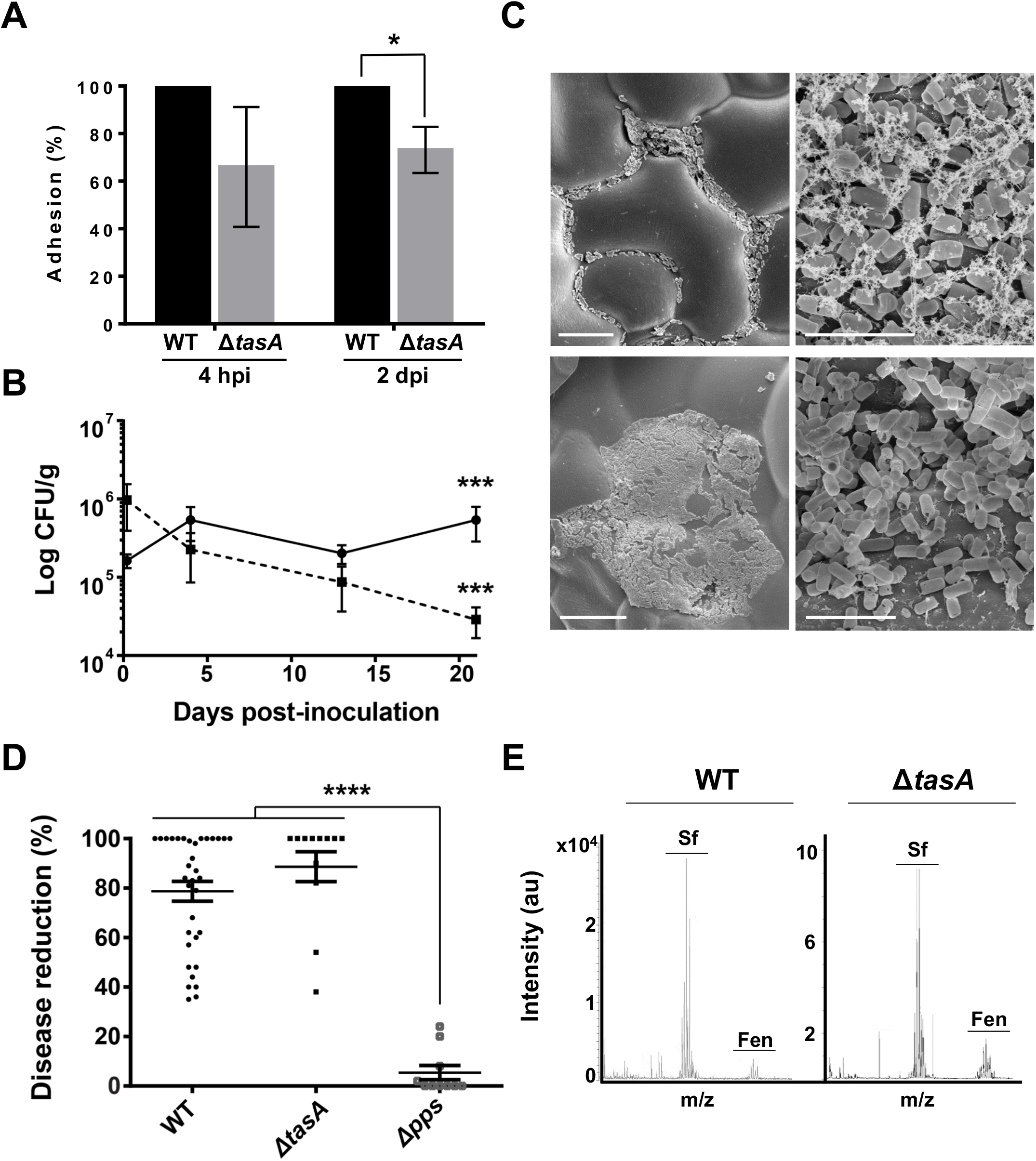
The amyloid protein TasA is essential for the fitness of *Bacillus* cells on the melon phylloplane. A) The adhesion of the WT (black bars) and *ΔtasA* (gray bars) strains to melon leaves at 4 h and 2 days post-inoculation showed statistically significant differences 2 days post-inoculation. Average values of three biological replicates are shown with error bars representing the SEM. Statistical significance was assessed via t-tests at each time-point (* p<0.05). B) Electron micrographs of inoculated plants taken 20 days post-inoculation show the WT cells (top) distributed in small groups covered by extracellular material and the *ΔtasA* cells (bottom) in randomly distributed plasters of cells with no visible extracellular matrix. Scale bars = 25 µm (left panels) and 5 µm (right panels) C) The persistence of the *ΔtasA* cells (dashed line) was significantly reduced compared with that of the WT cells (solid line). Average values of 5 biological replicates are shown with error bars representing the SEM. Statistical significance was assessed by independent t-test at each time-point (*** p<0.001). D) The WT and *ΔtasA* strains showed comparable biocontrol activity against the fungal phytopathogen *Podosphaera xanthii*. D) LC-MS-MS analysis revealed a higher fengycin level on melon leaves treated with *ΔtasA* (right) cells compared with that on leaves treated with WT cells (left).

Based on the reduced fitness exhibited by the single ECM component mutant strains and their deficiencies in biofilm formation, we hypothesized that these strains may also be defective in their antagonistic interactions with *Podosphaera xanthi* (an important fungal biotrophic phytopathogen of crops^42^) on plant leaves. Strains with mutations in *eps* and *bslA* partially ameliorated the disease symptoms, although their phenotypes were not significantly different from those of the WT strain (Fig. S1D). However, contrary to our expectations, the *ΔtasA* strain retained similar antagonistic activity to that of the WT strain (Fig. 1D). The simplest explanation for this finding is that the antifungal activity exhibited by the *ΔtasA* cells is due to higher production of antifungal compounds. *In situ* mass spectrometry analysis revealed a consistently higher relative amount of the antifungal compound plipastatin (also known as fengycin, the primary antifungal compound produced by *B. subtilis*) on leaves treated with *ΔtasA* cells compared to those treated with WT cells (Fig. 1E). These observations argue in favor of the relevance of the ECM and specifically TasA in the colonization and survival of *B. subtilis* on the phylloplane and revealed the importance of this ECM structural component in the antagonistic activity of *Bacillus* on the phylloplane.

### Loss of TasA causes a global change in the physiological state of the bacterial cells

The increased fengycin production and the previously reported deregulation of the expression pattern of the *tapA* operon in a *ΔtasA* mutant strain^9^ led us to explore whether loss of *tasA* disrupts the genetic circuitry of the cells. We sequenced and analyzed the whole transcriptomes of *ΔtasA* and WT cells grown *in vitro* on MSgg agar plates, a chemically-defined medium specifically formulated to support biofilm formation. Total RNA was extracted from colonies grown for 72 h, a time-point at which the phenotypic differences were clearly visible (Fig. S2A). RNA-seq analysis (suppl. Tables 1 and 2) showed that deletion of *tasA* resulted in pleiotropic effects on the overall gene expression profile of this mutant (Fig. 2A, Fig. S3A), with more than 800 differentially expressed genes (299 induced and 520 repressed), and these gene expression changes could hypothetically be responsible for substantial physiological changes (Fig. S4). A closer look at the data allowed us to cluster the differentially expressed genes into well-known regulons (Fig. 2A). The *sigKN*, *sigG*, *gerR* and *gerE* regulators, which control the expression of genes related to sporulation, were repressed in the *ΔtasA* cells, consistent with the delayed sporulation defect previously reported in ECM mutants^9, 30^ (Fig. S5). In contrast, the expression levels of biofilm-related genes, including the *epsA-O*, and *tapA* operons, were higher in the *ΔtasA* cells compared to their expression levels in WT cells. We found higher expression of the *slrR* transcriptional regulator (Suppl. table 1), which could explain the induction of the ECM-related genes^27^.

**Figure 2.**
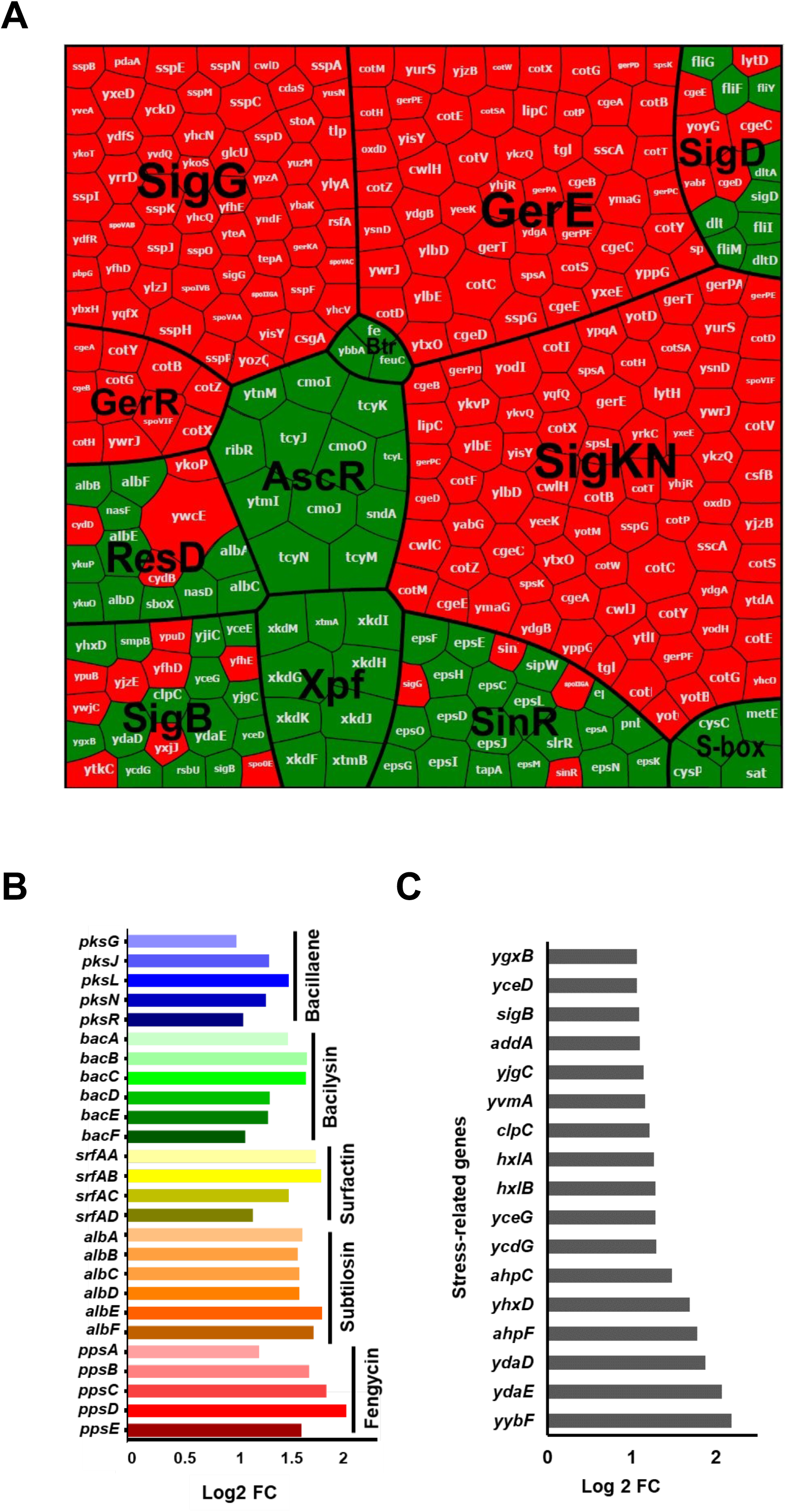
Whole-transcriptome analysis revealed major gene expression changes in the *ΔtasA* strain. A) Voronoi plot depicting the differentially expressed genes clustered into different regulons. The box size is proportional to the expression levels. Induced genes are colored in green, and repressed genes are colored in red. B) Genes related to the production of different secondary metabolites with antimicrobial activity that were induced in the transcriptomic analysis. C) Stress-related genes that were induced in the transcriptomic analysis.

An in-depth analysis of the transcriptional changes in the *ΔtasA* mutant cells highlighted the broad metabolic rearrangements that take place in *ΔtasA* colonies, including the induction of genes implicated in energy metabolism, secondary metabolism and general stress, among other categories (Suppl. table 1 and fig. S3A). First, the *alsS* and *alsD* genes, which encode acetolactate synthase and acetolactate decarboxylase, respectively, were clearly induced (Suppl. Table 1). This pathway feeds pyruvate into acetoin synthesis, a small four-carbon molecule that is produced in *B. subtilis* during fermentative and overflow metabolism^43^. Second, we observed induction of genes involved in fengycin biosynthesis (consistent with the overproduction of this antifungal lipopeptide *in planta*) (Fig. 1E), genes involved in the biosynthesis of surfactin, subtilosin, bacilysin and bacillaene (secondary metabolites with antimicrobial activities^44–47^) (Fig. 2B), as well as the operon encoding the iron-chelating protein bacillibactin (*dhbACEBF*) (Suppl. Table 1). All of these expression changes were confirmed via qRT-PCR analysis (Fig. S3B). Finally, the gene encoding the regulator AscR was induced. AscR controls transcription of the *snaA* (*snaAtcyJKLMNcmoOcmoJIrbfKsndAytnM)* and *yxe* (*yxeKsnaByxeMyxeNyxeOsndByxeQ*) operons (Suppl. Table 1), the products of which are members of alternative metabolic pathways that process modified versions of the amino acid cysteine. More specifically, the products of the *snaA* operon degrade alkylated forms of cysteine that are produced during normal metabolic reactions due to aging of the molecular machinery^26^. The *yxe* operon is implicated in the detoxification of S-(2-succino)cysteine, a toxic form of cysteine that is produced via spontaneous reactions between fumarate and cellular thiol groups in the presence of excess nutrients, which subsequently leads to increased bacterial stress^48, 49^. All of these transcriptional changes suggest an excess of cellular stress in the *ΔtasA* cells at 72 hours. In support of this prediction, an additional sign of stress was the overexpression of the sigma factor SigB (σ^B^) (suppl. Table 1), which controls the transcription of genes related to the general stress response^23^ (Fig. 2C), those encoding antibiotic resistance proteins and multidrug transporters (*ybbF* and *yvmA*), and proteins that confer resistance to other stressors, such as ethanol (*ydaD*, *yhxD*, *yceG*) or peroxide radicals (*ahpC* and *ahpF*) (Suppl. Table 1 and Fig. S3D). Interestingly, and also related to bacterial cell stress, we observed that nearly 67% of the genes belonging to the lysogenic bacteriophage PBSX were induced, a feature that has been reported to occur in response to mutations as well as to DNA or peptidoglycan damage^50, 51^ (suppl. Table 1).

In general, the transcriptional changes in the *ΔtasA* cells illustrate an intrinsic major physiological change suggesting excessive cellular stress and the early entry of the cells into stationary phase, which is supported by increased expression levels of genes related to: i) biofilm formation ii) synthesis of secondary metabolites (siderophores, antimicrobials, etc.); iii) fermentative metabolic pathways and overflow metabolism; iv) paralogous metabolism and assimilation of modified or toxic metabolic intermediates; v) general stress; and vi) induction of the lysogenic bacteriophage PBSX.

### *ΔtasA* cells exhibit low primary metabolic activity and increased secondary metabolism

Our transcriptomic analysis suggested that *ΔtasA* cells exhibit a shift from aerobic respiration to fermentation and anaerobic respiration as well as activation of secondary metabolism, physiological features typical of stationary phase cells^22, 52^. Activation of secondary metabolism provides an efficient means to utilize molecular intermediates upon growth arrest, nutrients depletion, or population stabilization. Fengycin is among the molecules produced during the later stages of bacterial growth. Based on the higher abundance of this molecule found on leaves treated with *ΔtasA* cells and its key role in the interaction between *B. subtilis* and fungal pathogens, we further investigated the kinetics of fengycin production *in vitro*. Flow cytometry analysis of cells expressing YFP under the control of the fengycin operon promoter demonstrated that fengycin production was induced in a subpopulation of cells (26.5%) at 48 h in the WT strain, reminiscent of the expression pattern reported for surfactin^41^. However, more than half of the *ΔtasA* population (67.3%) actively expressed YFP from the fengycin operon promoter at this time point (Fig. 3A top). At later stages of growth (72 h), the promoter was still active in the *ΔtasA* cells, and the population of positive cells was consistently higher than that in the WT strain (Fig. 3A bottom). Mass spectrometry analysis of cell-free supernatants demonstrated that this expression level was sufficient for the *tasA* mutant cells to produce nearly an order of magnitude more fengycin (Fig. 3B), consistent with our findings in plants (Fig. 1E). The fengycin levels found in the cell-free *ΔtasA* supernatant should be sufficient to provide at least the same level of antifungal activity as that extracted from the WT cells. *In vitro* experiments validated this hypothesis, showing that the cell-free supernatants from *ΔtasA* cells exhibited antifungal activity against *P. xanthii* conidia equivalent to that of WT cells, even in highly diluted spent medium (Fig. 3C). These results confirm the robust antimicrobial potency of *ΔtasA* cells and imply that primary metabolic intermediates are diverted to different pathways to support the higher secondary metabolite production in the *ΔtasA* mutant cells.

**Figure 3.**
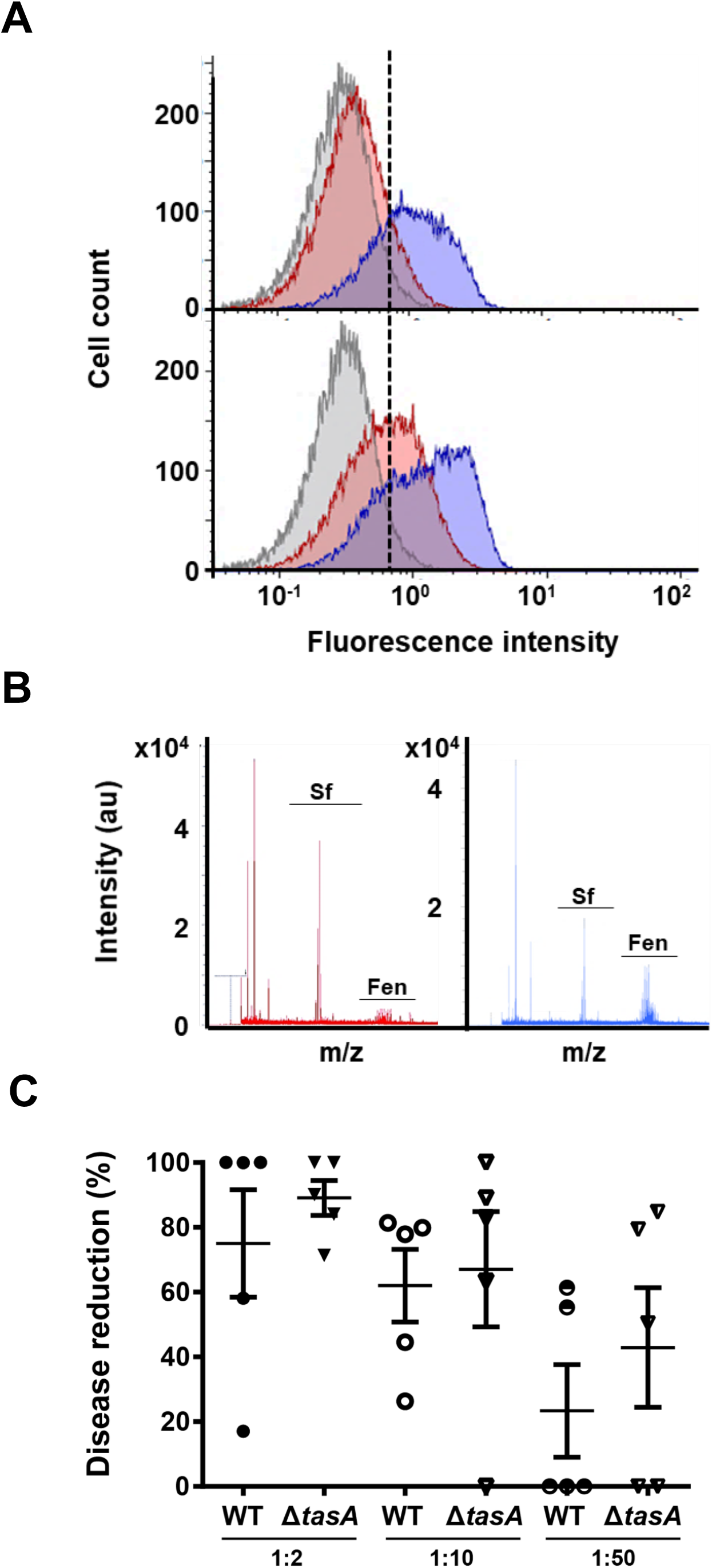
*ΔtasA* cells produce larger amounts of fengycin. A) Flow cytometry results with cells encoding the promoter of the fengycin promoter fused to YFP show that a higher percentage of *ΔtasA* cells (blue) expressed YFP compared with the percentage of YFP-expressing WT cells (red) at 48 h (top) and 72 h (bottom). B) MALDI-MS analysis of spent medium showed a higher fengycin level in *ΔtasA* cultures (right) compared to that in WT cultures (left). C) Serial dilutions of spent medium after 72 h of incubation showed that the medium from *ΔtasA* cultures retained as much antifungal activity as the medium from WT cultures. Average values of five biological replicates are shown. Error bars represent the SEM.

Consistent with these findings, we observed two complementary results that indicate less efficient metabolic activity in *ΔtasA* cells compared to that in WT cells. First, the *nasD* and *nasF* genes (parts of the anaerobic respiration machinery) were induced, and genes encoding several terminal oxidases found in the electron transport chain (*ythA*, *qoxB*, *cydD* and *cydB*) were differentially expressed (suppl. Table 1). The analysis of the respiration rates of these strains using the tetrazolium-derived dye 5-cyano-2,3-ditolyl tetrazolium chloride (CTC) and flow cytometry revealed a higher proportion of *ΔtasA* cells with lower respiration rates at 24 and 72 h compared to the WT proportions (69.10% vs. 43.07% at 24 h and 74.56 vs. 65.11% at 72 h, respectively) (Suppl. Table 3 and Fig. 4A). Second, the expression levels of the *alsSD* genes, which are responsible for the synthesis of acetoin (a metabolite produced by fermentative pathways) were higher in the *ΔtasA* strain than in the WT strain (Suppl. Table 1 and Fig. S3C). Indeed, all of the factors required for acetoin synthesis from pyruvate were overexpressed, whereas some key factors involved in the divergent or gluconeogenetic pathways were repressed (Suppl. Table 1 and Fig. S3E). Expression of *alsS* and *alsD* is induced by acetate, low pH and anaerobiosis^43, 53, 54^. Acetoin, in contrast to acetate, is a neutral metabolite produced to manage the intracellular pH and to ameliorate over-acidification caused by the accumulation of toxic concentrations of acetate or lactate, and its production is favored during bacterial growth under aerobic conditions^55^. Reduced respiration rates typically result in the accumulation of higher cellular proton concentrations, which leads to cytoplasmic acidification. These observations led us to postulate that the activation of the *alsSD* genes and the lower respiration rates observed in *ΔtasA* colonies might also reflect acidification of the intracellular environment, a potential cause of stationary phase-related stress. Measurements of the intracellular pH levels using the fluorescent probe 5-(6)carboxyfluorescein diacetate succinimidyl ester confirmed a significant decrease in the intracellular pH of nearly one unit (−0.92 ± 0.33, p value = 0.016) in *ΔtasA* cells at 72 h (Fig. 4B) compared to that in WT cells.

**Figure 4.**
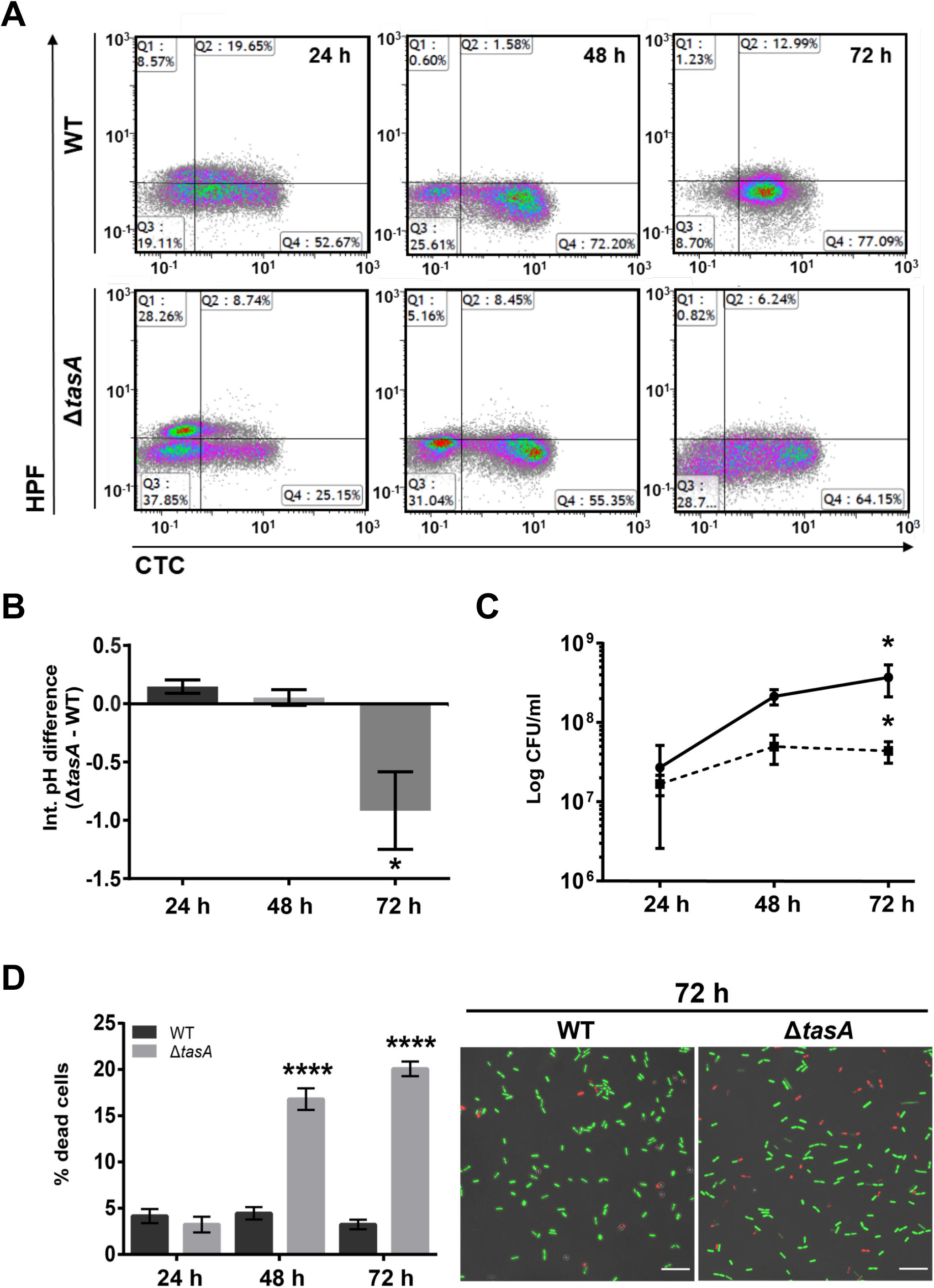
*ΔtasA* cells show altered respiration rates, lower intracellular pH, and increased cell death. A) Flow cytometry density plots of cells double stained with the HPF (Y axis) and CTC (X axis) dyes show that *ΔtasA* cells were metabolically less active (lower proportion of cells reducing CTC) and were under oxidative stress as early as 24 h (higher proportion of HPF-stained cells). B) Measurements of intracellular pH show significant cytoplasmic acidification in the *ΔtasA* cells at 72 h. Average values of four biological replicates are shown. Statistical significance was assessed by one-way ANOVA with the Tukey test (* p<0.05). C) The population dynamics in *ΔtasA* (dashed line) and WT colonies (solid line) grown on MSgg agar at 30 °C showed a difference of nearly one order of magnitude in the *ΔtasA* colony from 48 h. Average values of three biological replicates are shown. Error bars represent the SEM. Statistical significance was assessed by the Mann-Whitney test (* p<0.05. D) Left. Quantification of the proportion of dead cells treated with the BacLight LIVE/DEAD viability stain in WT and *ΔtasA* colonies at different time-points reveled a higher population of dead cells in *ΔtasA* colonies compared to that found in the WT colonies. Average values of three biological replicates are shown with error bars representing the SEM. For each experiment, at least three fields-of-view were measured. Statistical significance was assessed via independent t-tests at each time-point (**** p<0.0001). Right. Representative confocal microscopy images of fields corresponding to LIVE/DEAD stained WT or *ΔtasA* samples at 72 h. Scale bars = 10 µm.

Based on these results, we conclude that the *ΔtasA* mutant presents alterations in primary and secondary metabolism, the latter of which were more robust, that lead to overproduction of secondary metabolites and over-acidification of the intracellular environment.

### Loss of TasA increases membrane fluidity and cell death

The reduction in metabolic activity of *ΔtasA* cells, along with their acidification of the intracellular environment, might be expected to result in reduced bacterial viability. Measurements of the dynamics of viable bacterial cell density, expressed as CFU counts, showed that after 48 h, *ΔtasA* colonies possessed nearly an order of magnitude fewer CFUs than did WT colonies (Fig. 4C). These results suggest the hypothesis that *ΔtasA* colonies might exhibit higher rates of cell death than WT colonies. To validate this hypothesis, we analyzed the live and dead sub-populations using the BacLight LIVE/DEAD viability stain and confocal microscopy (Fig. 4D left). The proportion of dead cells in *ΔtasA* colonies ranged from between 16.80% (16.80 ± 1.17) and 20.06% (20.06 ± 0.79) compared to 4.45% (4.45 ± 0.67) and 3.24% (3.24 ± 0.51) found in WT colonies at 48 and 72 h, respectively (Fig. 4D right). The significantly higher rate of cell death in *ΔtasA* compared to WT is consistent with the drastically lower bacterial counts found in the *ΔtasA* mutant colonies after 48 hours.

Apoptosis or programmed cell death (PCD) is a well-established process in eukaryotic cells whose existence in prokaryotes has recently been accepted^56–58^. One of the instigators of PCD is the age-dependent accumulation of damaged cellular components and flaws in metabolic reactions. The accumulation of reactive oxygen species (ROS) is a well-known trigger of PCD^59^. To determine if *ΔtasA* cells possess abnormal ROS levels, we monitored ROS generation using hydroxyphenyl fluorescein (HPF), a fluorescent indicator of the presence of highly reactive hydroxyl radicals. Flow cytometry analysis revealed a larger proportion of HPF-positive cells (which have increased ROS levels) in the *ΔtasA* strain at 24 h compared to the WT proportion (42.38% vs. 28.61%, respectively) (Fig. 4A and suppl. table 3). ROS are highly reactive, unstable molecules that target different cellular components, including DNA and lipid bilayers. Indeed, DNA damage and ion gradient disruption (which translates into altered membrane potential) are two hallmarks of PCD in both prokaryotes and eukaryotes. We first searched for DNA damage by performing TUNEL assays to fluorescently stain bacterial cells containing DNA strand breaks. At 24 and 48 hours, we found a significantly higher number of fluorescently-stained *ΔtasA* cells compared with the number of fluorescently-stained WT cells (shown in Fig. 5A and quantified in 5B). These results indicated that DNA damage appears to occur not only earlier, but also with a higher frequency, in *tasA* cells than in WT cells. A sizeable number of stained cells was also found at 72 h in the *ΔtasA* colonies, the same time-point at which the TUNEL signal started to increase in the WT colonies (Fig. 5A). The TUNEL signal in the *ΔtasA* cells at this time-point was not significantly different from that of the WT cells (Fig. 5B), probably due to the increased cell death in the *ΔtasA* cells.

**Figure 5.**
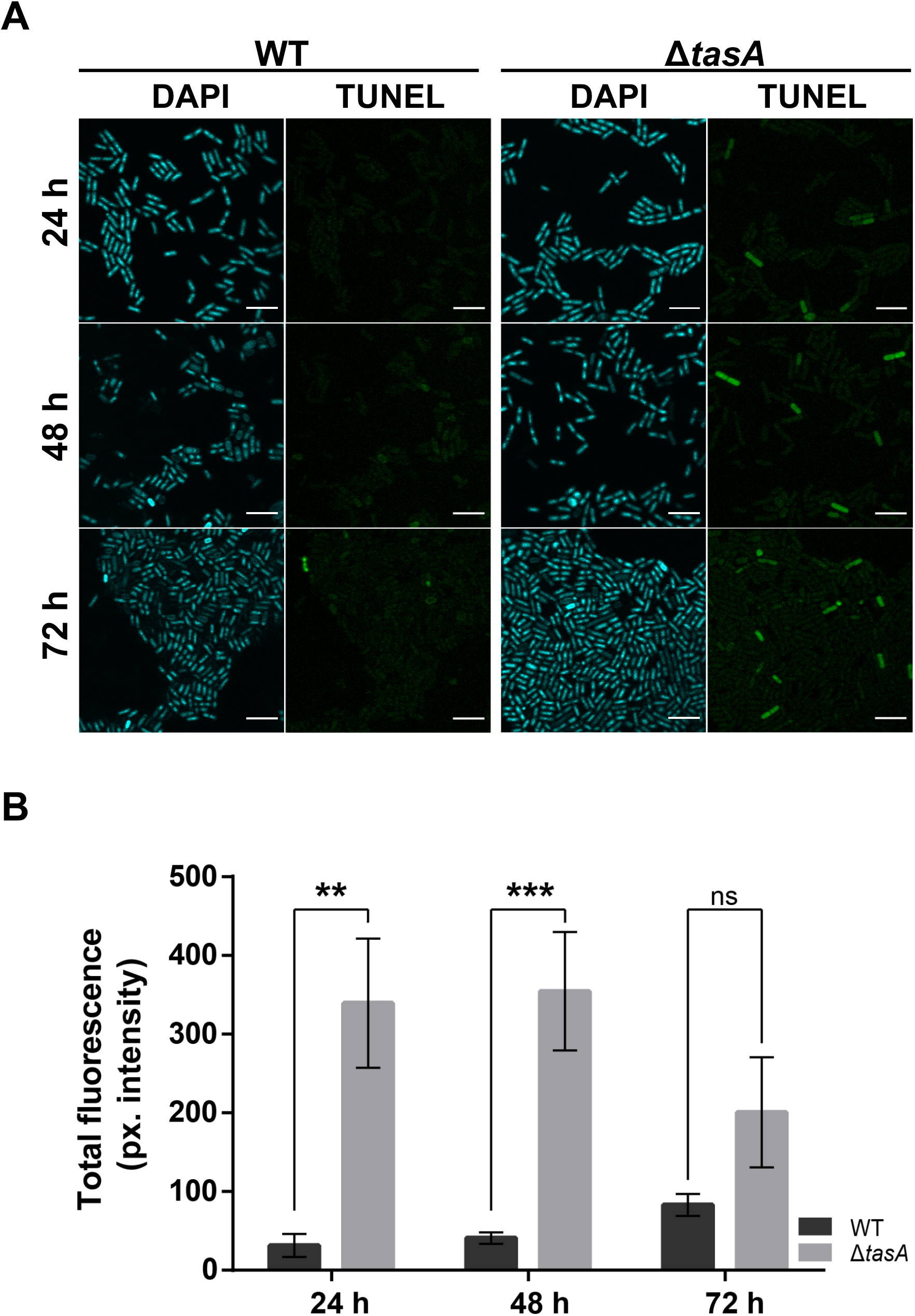
The *ΔtasA* cells exhibit higher levels of DNA damage. A) TUNEL assays (right) and CLSM analysis revealed significant DNA damage in the *ΔtasA* cells compared to that in the WT cells. The cells were counterstained with DAPI DNA stain (left). Scale bars = 5 µm. B) Quantification of the TUNEL signals in WT and *ΔtasA* colonies. The results showed significant differences in the DNA damage levels in *ΔtasA* and WT cells after 24 and 48 h of growth. Average values of three biological replicates are shown. For each experiment, at least three fields-of-view were measured. Error bars indicate the SEM. Statistical significance was assessed via independent t-tests at each time-point (** p<0.01 *** p<0.001)

Next, we assessed the cellular membrane potential using the fluorescent indicator tetramethylrhodamine, methyl ester (TMRM). Consistent with the levels of DNA damage detected via the TUNEL assays, the alterations in the membrane potential of the *ΔtasA* cells were significantly different at all time points compared with the corresponding values for the WT cells (Fig. 6A left panel and 6B left). These results indicate that after 48 h (the same time point at which the cell death rate increases and the cell population plateaus in *ΔtasA* colonies) *ΔtasA* cells also exhibit increased membrane hyperpolarization compared with that in the WT cells, a feature that has been linked to mitochondrial-triggered PCD in eukaryotic cells^60–62^.

**Figure 6.**
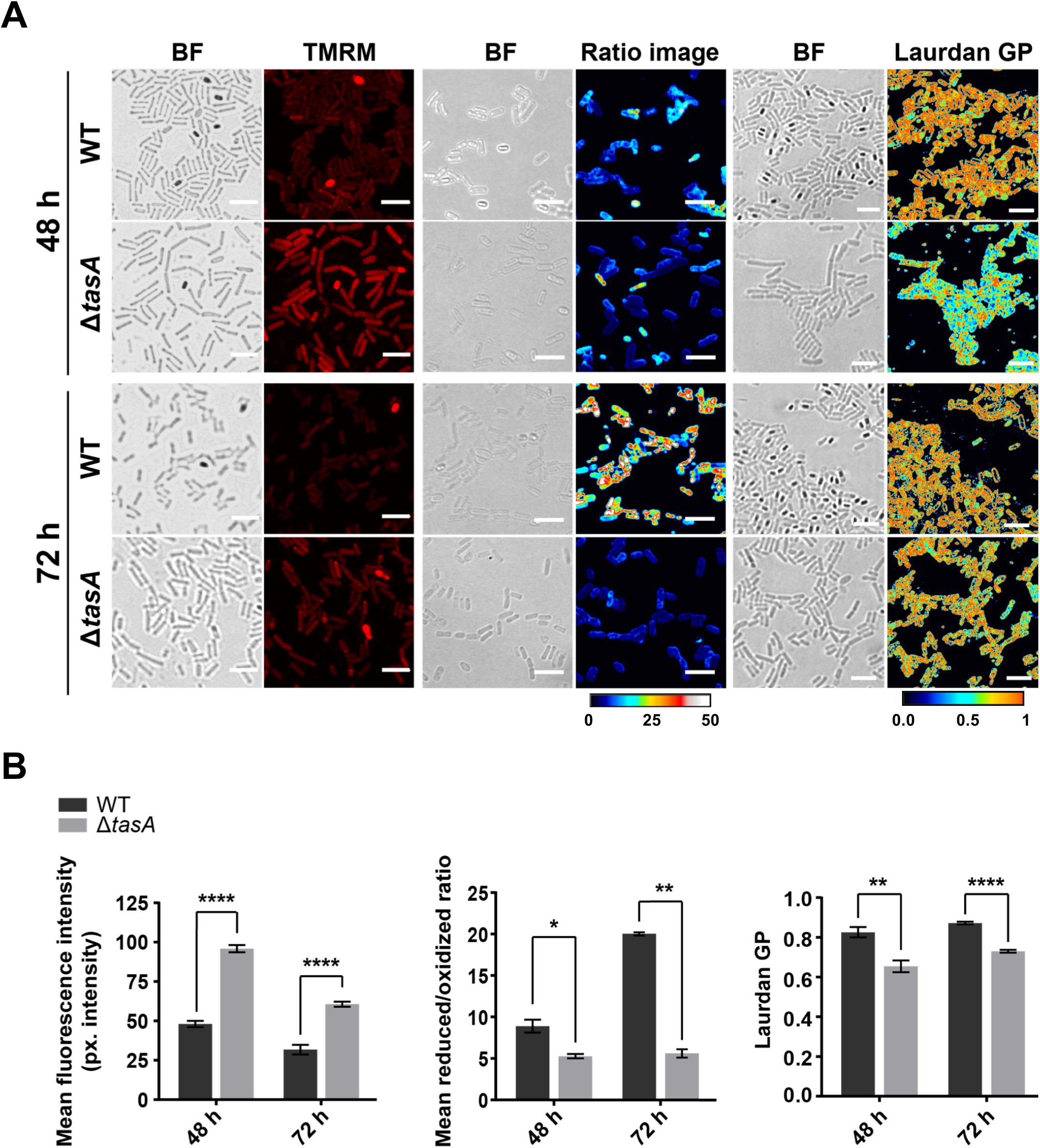
*ΔtasA* cells show altered membrane potential, higher susceptibility to lipid peroxidation and exhibit high membrane fluidity. A) Left panel. A TMRM assay of WT and *ΔtasA* cells, located at the top or bottom respectively in each set, showed a decrease in membrane potential in the WT cells, whereas the *ΔtasA* cells exhibited hyperpolarization at 48 and 72 h. Center panel. Assessment of the lipid peroxidation levels using BODIPY 581/591 C11 reagent in WT and *ΔtasA* cells after treatment with 5mM CuHpx and analysis by CLSM. The ratio images represent the ratio between the two states of the lipid peroxidation sensor: reduced channel_(590-613 nm emission)_/oxidized channel_(509-561nm emission)_. The ratio images were pseudo-colored depending on the reduced/oxidized ratio values. A calibration bar (from 0 to 50) is located at the bottom of the panel. Confocal microscopy images show that CuHpx treatment was ineffective in the WT strain at 72 h, whereas the mutant strain showed symptoms of lipid peroxidation. Right panel. Laurdan GP analyzed via fluorescence microscopy. The images were taken at two different emission wavelengths (gel phase, 432 to 482 nm and liquid phase, 509 to 547 nm) that correspond to the two possible states of the Laurdan reagent depending on the lipid environment. The Laurdan GP images represent the Laurdan GP value of each pixel (see Materials and methods). The Laurdan GP images were pseudo-colored depending on the laurdan GP values. A calibration bar (from 0 to 1) is located at the bottom of the set. The Laurdan GP images show an increase in membrane fluidity (lower Laurdan GP values) in the *tasA* mutant cells 48 and 72 h. All scale bars are equal to 5 µm. B) Quantification of the TMRM signal, lipid peroxidation and laurdan GP revealed statistically significant differences between the WT and *ΔtasA* colonies at 48 and 72 h. Average values of three biological replicates are shown with error bars representing the SEM. For each experiment, at least three fields-of-view were measured. Statistical significance in each experiment was assessed via independent t-tests at each time-point (**** p<0.0001, * p<0.05, ** p<0.01).

The differences in ROS production, DNA damage level and membrane hyperpolarization between the WT and *ΔtasA* cell populations are consistent with increased PCD being the cause of the higher cell death rate observed in *ΔtasA* colonies after 24 h. To test the idea that loss of *tasA* results in increased PCD, we investigated the level of membrane lipid peroxidation, a chemical modification derived from oxidative stress that subsequently affects cell viability by inducing toxicity and apoptosis in eukaryotic cells^63, 64^. Staining with BODIPY 581/591 C11, a fluorescent compound that is sensitive to the lipid oxidation state and localizes to the cell membrane, showed no significant detectable differences in the levels of lipid peroxidation at any time point (Fig. S11C). However, treatment with cumene hydroperoxide (CuHpx), a known inducer of lipid peroxide formation^65^, resulted in different responses in the two strains. WT cells showed high reduced/oxidized ratios at 48 and 72 h and, thus, a low level of lipid peroxidation (Fig. 6A center panel and fig. 6B center). In contrast, the comparatively lower reduced/oxidized ratios in *ΔtasA* cells at 48 and 72 h indicated increased lipid peroxidation (fig. 6A center panel and fig. 6B center). These results demonstrate that the *ΔtasA* strain is less tolerant to oxidative stress than is the WT strain, and, therefore, is more susceptible to ROS-induced damage.

Based on the higher susceptibility of the *ΔtasA* strain to lipid peroxidation and considering the differences in ROS production between the WT and *ΔtasA* cells, we next studied the integrity and functionality of the plasma membrane. First, no clear differences in the integrity, shape or thickness of the cell membrane or cell wall were observed via transmission electron microscopy (TEM) of negatively stained thin sections of embedded *ΔtasA* or WT cells at 24 and 72 h under our experimental conditions (Fig. S6). Next, we examined membrane fluidity, an important functional feature of biological membranes that affects their permeability and binding of membrane-associated proteins, by measuring the Laurdan generalized polarization (Laurdan GP)^63, 66^. Our results show that the Laurdan GP values were significantly lower at 48 and 72 h in *ΔtasA* cells compared with the values in WT cells (0.65 ± 0.03 or 0.82 ± 0.03 p value = 0.0001, respectively, at 48 h, and 0.87 ± 0.006 or 0.73 ± 0.007 p value < 0.0001, respectively, at 72 h) (Fig. 6A right panel and 6B right). These results indicate incremental changes in membrane fluidity, comparable to that resulting from treatment of cells with benzyl alcohol, a known membrane fluidifier (Fig. S13A top and center panels and fig. S13B). Membrane fluidity has been associated with higher ion, small molecule and proton permeability^67, 68^, which would explain why the *ΔtasA* cells are impaired in energy homeostasis as well as the subsequent effects on the intracellular pH and membrane potential that eventually contribute to cell death.

### TasA is located in the DRM of cell membranes

The negative effects on membrane potential and fluidity observed in the *ΔtasA* cells suggest alterations in membrane dynamics, which in bacterial cells are directly related to functional membrane microdomains (FMM); FMMs are specialized membrane domains that also regulate important cellular functions such as KinC-dependent biofilm formation, sporulation, protein secretion, competence, motility or cell division among others^69–72^. The bacterial flotillins FloT and FloA are localized in FMMs and are directly involved in the regulation of membrane fluidity^69^. Our transcriptomic data of *ΔtasA* colonies at 72 h revealed induction of genes that encode proteins that interact with either FloT alone or with FloT and FloA (suppl. table 4)^70^. These two lines of evidences led us to propose a connection between the membrane fluidity and permeability of *ΔtasA* cells and changes in the FMMs. We initially studied the membrane distribution of FloT as a marker for FMMs in WT and *ΔtasA* cells using a FloT-YFP translational fusion construct and confocal microscopy (Fig. 7A). The WT strain showed the typical FloT distribution pattern, in which the protein is located within the bacterial membrane in the form of discrete foci^73^ (Fig. 7A top). However, in the *ΔtasA* cells, the fluorescent signal was visible only in a subset of the population, and it was completely mislocalized (Fig. 7A bottom). In agreement with these findings, quantification of the fluorescent signal in WT and *ΔtasA* samples showed significant decreases in the signal in the *ΔtasA* mutant cells at 48 and 72 h (Fig. 7B).

**Figure 7.**
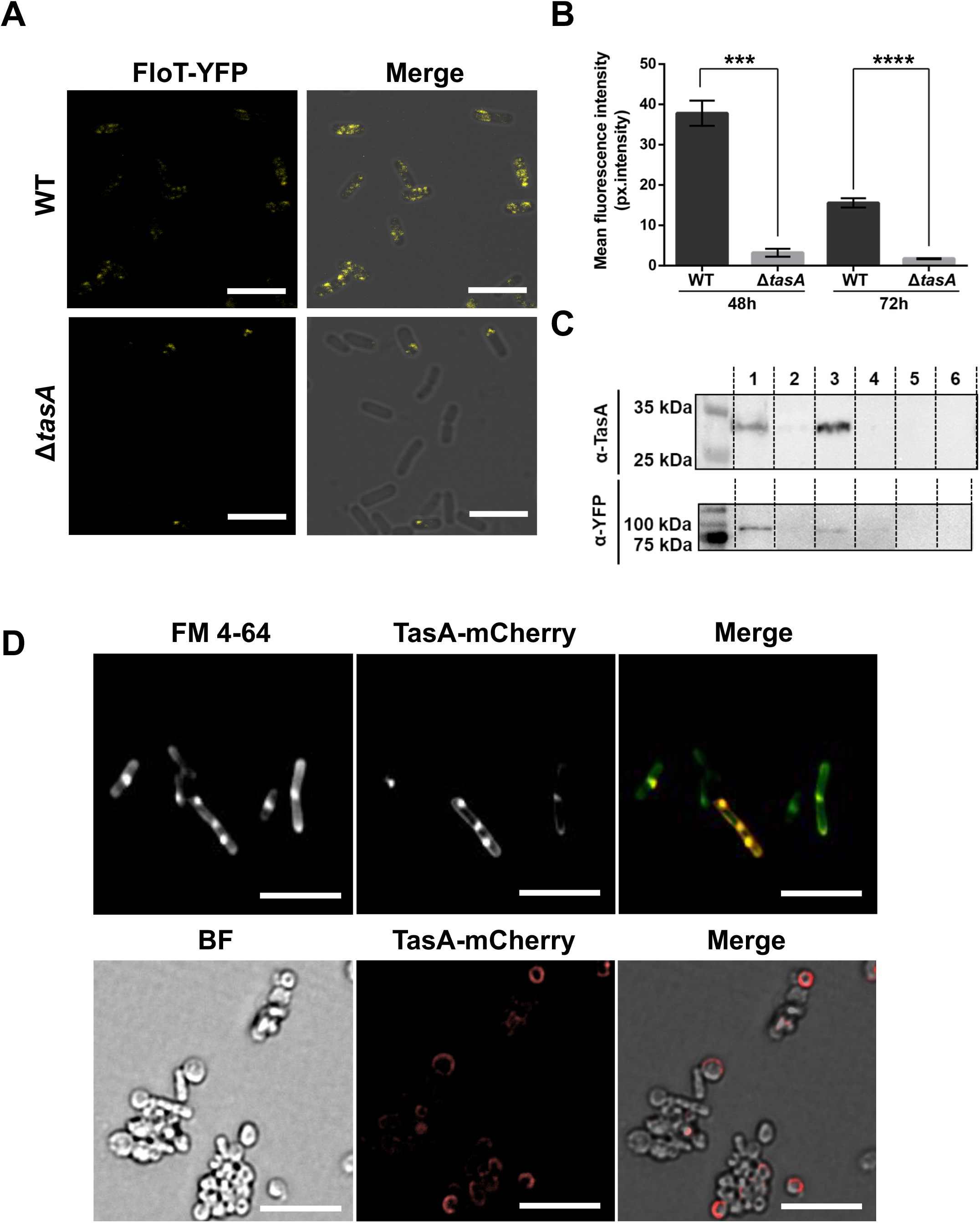
The absence of TasA causes mislocalization of the flotillin-like protein FloT. A) Representative confocal microscopy images showing WT or *ΔtasA* cells expressing the *floT-yfp* construct at 72 hours. WT images show the typical punctate pattern associated to FloT. That pattern is lost in *ΔtasA* cells. B) Quantification of fluorescence signal in WT and *ΔtasA* samples at 48 and 72 h show significant differences between the two strains (*** p<0.001 **** p<0.0001). Average values of three biological replicates are shown with error bars representing the SEM. For each experiment, at least three fields-of-view were measured. C) Western blot of different membrane fractions exposed to an anti-TasA or anti-YFP antibodies. 1 = WT DRM fraction, 2 = WT DSM fraction, 3 = WT cytosolic fraction, 4 = *ΔtasA* DRM fraction, 5 = *ΔtasA* DSM fraction, 6 = *ΔtasA* cytosolic fraction. D) Top. Fluorescence microscopy of 72 h WT cells expressing *tasA-mCherry* stained with the membrane dye FM 4-64. Bottom. Fluorescence microscopy of protoplast cells expressing a *tasA-mCherry* construct. The signal corresponding to TasA appears in the surface of protoplasts. Scale bars = 5

The different FloT localization observed in the *ΔtasA* mutant cells led us to speculate on the presence of TasA in FMMs. Membranes from both prokaryotes and eukaryotes can be separated into detergent-resistant (DRM) and detergent-sensitive fractions (DSM) based on their solubility in detergent solutions^73^. Although it is important to point out that the DRM and FMMs (or lipid rafts in eukaryotes) are not equivalent, the DRM fraction has a differential lipid composition and is enriched with proteins, rendering it more resistant to detergents; furthermore, many of the proteins present in FMMs are also present in the DRM^74^. Immunodetection assays of the DRM, DSM and cytosolic fractions of each strain using an anti-TasA antibody showed the presence of anti-TasA reactive bands of the expected size primarily in the DRM fraction and in the cytosol (Fig. 7C top, lanes 1 and 3). As controls, the fractions from the *tasA* mutant showed no signal (Fig. 7C top, lanes 4, 5 and 6). Western blots of the same fractions isolated from WT and *ΔtasA* strains carrying a FloT-YFP translational fusion with an anti-YFP antibody (Fig. 7C bottom) confirmed that FloT was mainly present in the DRM of WT cells (Fig. 7C bottom, lane 1). The signal was barely noticeable in the same fraction from *ΔtasA* cells (Fig. 7C bottom, lane 4), mirroring the reduced fluorescence levels observed via microscopy (Fig. 7A). We further validated the presence of TasA in the bacterial cell membrane by using a strain carrying a TasA-mCherry translational fusion construct. Fluorescence microscopy showed an overlap between the TasA-mCherry signal and the membrane-specific dye FM 4-64 (Fig. 7D, top). Second, the surfaces of protoplasted cells, i.e., cells lacking the peptidoglycan layer, were decorated with fluorescent signal corresponding to the TasA-mCherry fusion protein (Fig. 7D bottom). Based on these results, we concluded that TasA is located in the DRM fraction of the cell membrane and that its absence leads to mislocalization and loss of the flotillin-like protein FloT and alterations in membrane dynamics and fluidity.

### Mature TasA is required to maintain viable bacterial physiology

TasA is a secreted protein located in the cell membrane and ECM, and reaching these sites requires the aid of secretion-dedicated chaperones, the translocase machinery and the membrane-bound signal peptidase SipW^75^. It is known that TasA processing is required for assembly of the amyloid fibrils and biofilm formation^35, 76^. However, formation of the mature amyloid fibril requires the accessory protein TapA, which is also secreted via the same pathway^36^, is present in the mature amyloid fibers and is found on the cell surface^76^. Considering these points, we first wondered whether TapA is involved in the increased PCD observed in the *ΔtasA* mutant. By applying the BacLight LIVE/DEAD viability stain to a *ΔtapA* colony, we found a similar proportion of live to dead cells as that found in the WT colony at 72 h (Fig. 8A). *ΔtapA* cells lack TasA fibers but still expose TasA in their surfaces^76^; thus we reasoned that mature TasA is necessary for preserving the cell viability levels observed in the WT strain. To test this possibility, we constructed a strain bearing a mutation in the part of the *<u>tasA</u>* gene that encodes the TasA signal peptide ^77^. To avoid confounding effects due to expression of the mutated *tasA* gene in the presence of the endogenous operon, we performed this analysis in a strain in which the entire *tapA* operon was deleted and in which the modified operon encoding the mutated *tasA* allele was inserted into the neutral *lacA* locus. The strain carrying this construct was designated as “TasA SiPmutant”. The endogenous version of TasA successfully restored biofilm formation (Fig. S2B), while the phenotype of SiP mutant on MSgg medium at 72 h was different from those of both the WT and *tasA* mutant strains (Fig. 8B and Fig. S2B). Immunodetection analysis of TasA in fractionated biofilms confirmed the presence of TasA in the cells and ECM fractions from the WT strain and the strain expressing the endogenous version of *tasA* (Fig. 8C lanes 1 and 2 and 4 and 5 respectively). However, a faint anti-TasA reactive signal was observed in both fractions of the SiP mutant (Fig. 8C lanes 3 and 6). This result indicates that TasA is not efficiently processed in the SiP mutant and, thus, the protein levels in the ECM were drastically lower. The faint signal detected in the cell fraction might be due to the fact that the pre-processed protein is unstable in the cytoplasm and is eventually degraded over time^77^. Consistent with our hypothesis, the levels of cell death in the SiP mutant were significantly different from those of the WT strain (Fig. 8D). Taken together, these results rest relevance to TapA to the increase PCD observed in the absence of TasA and indicate that TasA must be processed to preserve the level of cell viability found in WT colonies.

**Figure 8.**
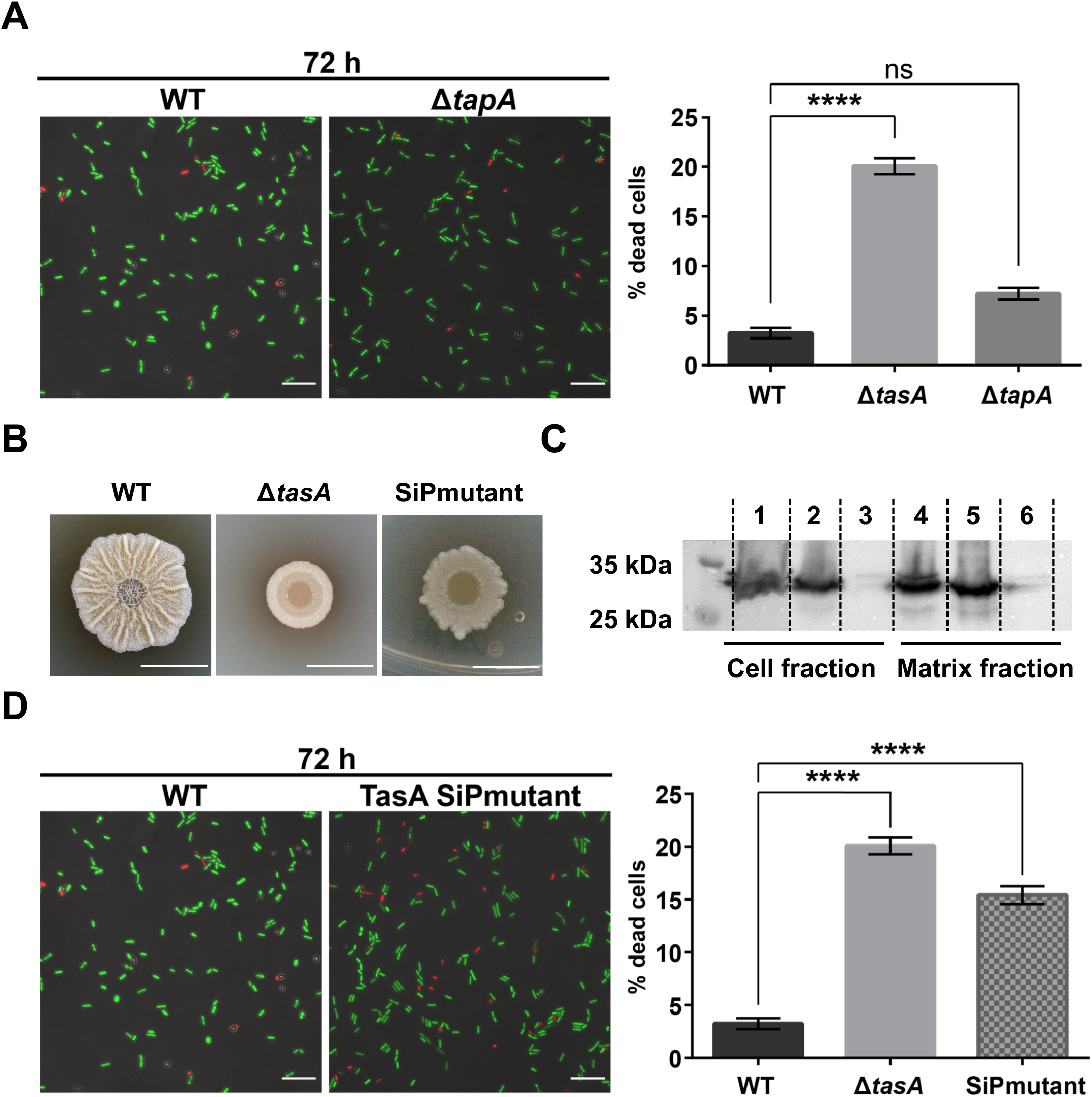
Mature TasA is required to stabilize the levels of cell death within the *B. subtilis* colony. A) Left. Representative confocal microscopy images of fields corresponding to LIVE/DEAD-stained WT or *ΔtapA* samples at 72 h. Scale bars = 10 µm. Right. Quantification of the proportion of dead cells in WT and *ΔtapA* colonies at 72 h reveled no differences in the levels of cell death. Average values of three biological replicates are shown with error bars representing the SEM. For each experiment, at least three fields-of-view were measured. Statistical significance was assessed via independent t-tests at each time-point (**** p<0.0001). B) Colony phenotypes of WT, *ΔtasA* and SiPmutant strains on MSgg agar 72 h. Scale bars = 1 cm. C) Western blot of the cell and matrix fractions of the three strains at 72 h exposed to an anti-TasA antibody. Lanes: 1 = WT cell fraction, 2 = *tasA*_native_ cell fraction 3 = SiPmutant cell fraction 4 = WT matrix fraction 5 = *tasA*_native_ matrix fraction 6 = SiPmutant matrix fraction. D) Left. Representative confocal microscopy images of fields corresponding to LIVE/DEAD stained WT or SiPmutant samples at 72 h. Scale bars = 10 µm. Right. Quantification of the proportion of dead cells in WT and SiPmutant colonies at different time points reveled higher levels of cell death in the SiPmutant colonies. Average values of three biological replicates are shown with error bars representing the SEM. For each experiment, at least three fields-of-view were measured. Statistical significance was assessed via independent t-tests at each time-point (**** p<0.0001).

### Cells expressing a TasA variant have a restored physiological status but are impaired in biofilm formation

The fact that the *ΔtapA* strain does not form TasA fibers but does have normal cell death as well as the increased membrane fluidity and mislocalization of the flotillin-like protein FloT in the *ΔtasA* strain led us to hypothesize that the TasA in the DRM, and not that in the ECM, is responsible for maintaining the PCD levels within the WT colonies. To validate this hypothesis, we performed an alanine scanning experiment with TasA to obtain an allele encoding a stable version of the protein that could support biofilm formation. To produce these constructs, we used the same genetic background described in the above section. The strain JC81, which expresses the allele TasA_p._[_Lys68_Asp69delinsAlaAla_], failed to rescue the biofilm formation phenotype in the WT strain (Fig. 9A and Fig. S7A). Immunodetection analysis of TasA in fractionated biofilms confirmed the presence of the mutated protein in the cells and in the ECM, indicating a malfunction in the protein’s structural role in proper ECM assembly (Fig. 9B lanes 1-4 and S8B). The cell membrane fractionation analysis revealed, however, the presence of mutated TasA in the DRM, DSM and cytosolic fractions (Fig. 9B lanes 5-7). Accordingly, JC81 was reverted to a physiological status comparable to that of the WT strain. This feature was demonstrated by similar expression levels of genes encoding factors involved in the production of secondary metabolites (i.e., *ppsD*, *albE*, *bacB*, *srfAA*) or acetoin (*alsS*), indicating comparable metabolic activities between the two strains (Fig. 9C). Further evidence confirmed the restoration of the metabolic status in JC81. First, similar proportions of WT and JC81 cells expressing YFP from the fengycin operon promoter were detected after 72 h of growth via flow cytometry analysis (Fig. 9D, green curve). In agreement with these findings, there were no differences in the proportions of cells respiring or accumulating ROS or in the intracellular pH values between the JC81 and WT strains (Fig. 9E and Fig. 9F). Consistently, the population dynamics of JC81 resembled that of the WT strain (fig. 9G), and, as expected, its level of cell death was comparable to that of the WT strain (Fig. 9H). Finally, there were no differences in any of the examined parameters related to oxidative damage and PCD (i.e., DNA damage, membrane potential, susceptibility to lipid peroxidation and membrane fluidity) between JC81 and WT cells (Fig. S8, Fig. S9, Fig. S11, and Fig. S12, respectively). Taken together, these findings assign TasA complementary functions in the stabilization of cell membrane dynamics and cellular physiology during normal colony development that prevent premature PCD, a role beyond the well-known structural function of amyloid proteins in biofilm ECMs.

**Figure 9.**
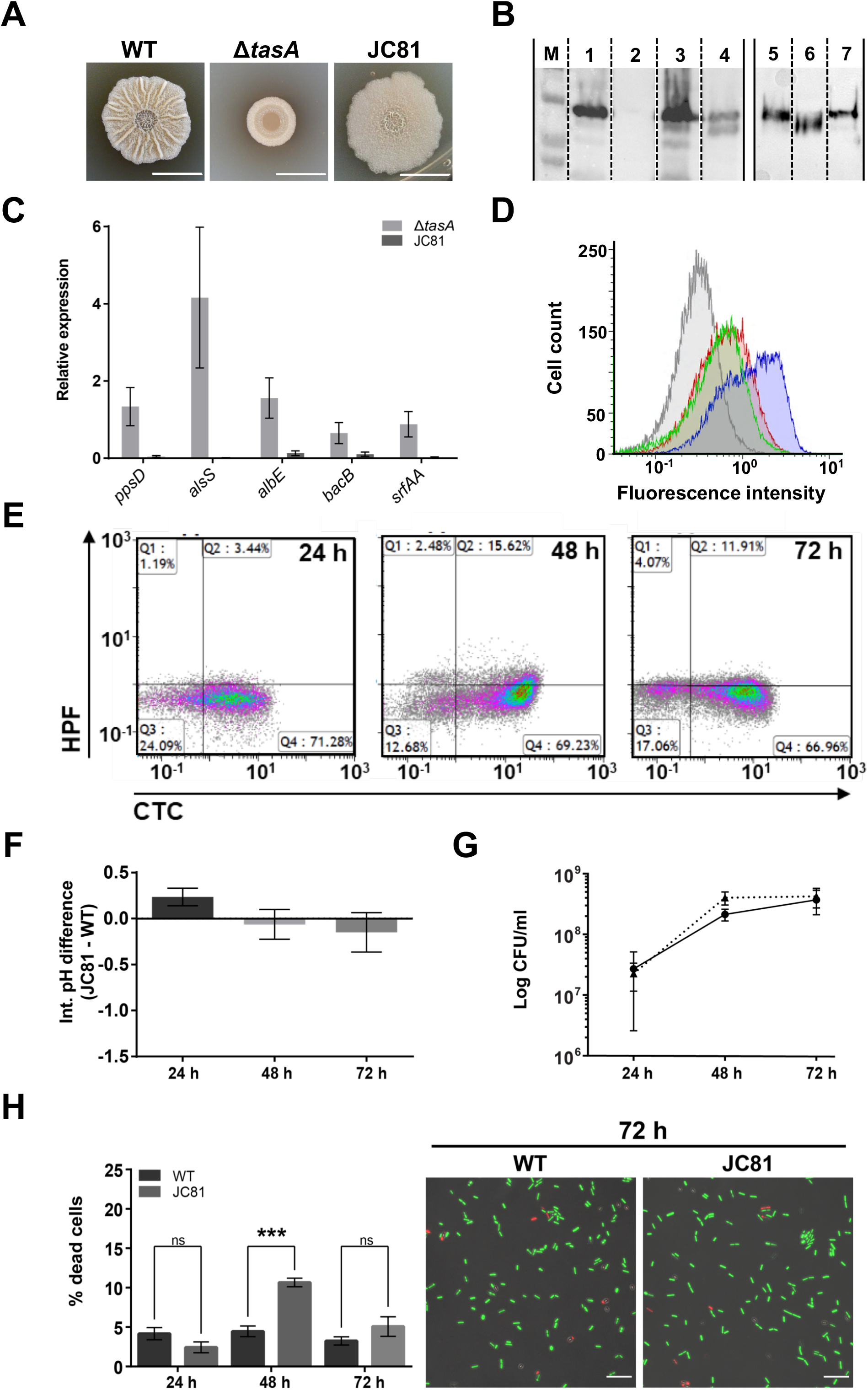
A TasA variant fails to restore biofilm formation in *tasA-*deleted cells, but rescues their physiological status. A) Colony phenotypes of the three strains on MSgg agar 72 h. Scale bars = 1 cm. B) A western blot of the cell fractions of the different strains at 72 h exposed to an anti-TasA antibody showed that the protein is present in JC81. Lanes: 1 = WT, 2 = *ΔtasA*, 3 = JC70 (*tasA*_native_) 4 = JC81 (*tasA*_variant_) 5 = DRM JC81 6 = DSM JC81 7 = cytosol JC81. C) The relative expression levels of *ppsD*, *alsS*, *albE*, *bacB* and *srfAA* in JC81 are similar to those in the WT strain. (dark gray bars = *ΔtasA*, light gray bars = JC81). Average values of three biological replicates are shown with error bars representing the SEM. D) Flow cytometry showed that the proportion of cells transcribing from the fengycin promoter in JC81 (green curve) was similar to that in the WT strain (red curve). Blue curve = *ΔtasA*. E) Density plots of cells double stained with the HPF (Y axis) and CTC (X axis) dyes show that JC81 behaved similarly to the WT strain. F) Intracellular pH measurements showed no difference between the WT and JC81 strains. Average values of four biological replicates are shown. Error bars represent the SEM. G) The population dynamics, in terms of CFU counts, in the JC81 colonies were similar to those in the WT colonies. Average values of four biological replicates are shown. Error bars represent the SEM. H) Quantification of the proportion of dead cells treated with the BacLight LIVE/DEAD viability stain in WT and JC81 colonies at different time-points. Average values of three biological replicates are shown with error bars representing the SEM. For each experiment, at least three fields-of-view were measured. Statistical significance was assessed via independent t-tests at each time-point (*** p<0.001). Representative confocal microscopy images of fields corresponding to LIVE/DEAD-stained WT or JC81 samples at 72 hours. Scale bars = 10 µm.

### The TasA variant impairs *B. subtilis* survival and fitness on the phylloplane

Our analysis of the intrinsic physiological changes in *ΔtasA* cells showed how the absence of TasA leads to the accumulation of canonical signs of PCD, a physiological condition typical of stationary phase cells, and, ultimately, to the premature aging of bacterial colonies. These observations help to reconcile two *a priori* contradictory features of *B. subtilis* ecology on plant leaves: the reduced persistence of the *ΔtasA* mutant on the melon phylloplane versus its ability to efficiently exert biocontrol against the fungus *P. xanthi*, which occurs via overproduction of fengycin and other antimicrobial molecules. Following this line of thought, we predicted that the JC81 strain, which expresses a version of TasA that is unable to restore biofilm formation but preserves the physiological status of the cells, would show overall signs of reduced fitness on melon leaves. The JC81 cells retained their initial ability to adhere to melon leaves (Fig. 10A); however, their persistence decreased (Fig. 10B) and their colonization showed a pattern somewhat intermediate between those of the WT and *ΔtasA* strains (Fig. 10C). In agreement with our prediction, the reduced fitness of this strain resulted in a failure to manage *P. xanthi* infection (Fig. 10D). Thus, we conclude that the ECM, by means of the amyloid protein TasA, is required for normal colonization and persistence of *B. subtilis* on the phyllosphere. These ecological features depend on at least two complementary roles of TasA: one role related to ECM assembly and a new proposed role in the preservation of the physiological status of cells via stabilization of membrane dynamics and the prevention of PCD.

**Figure 10.**
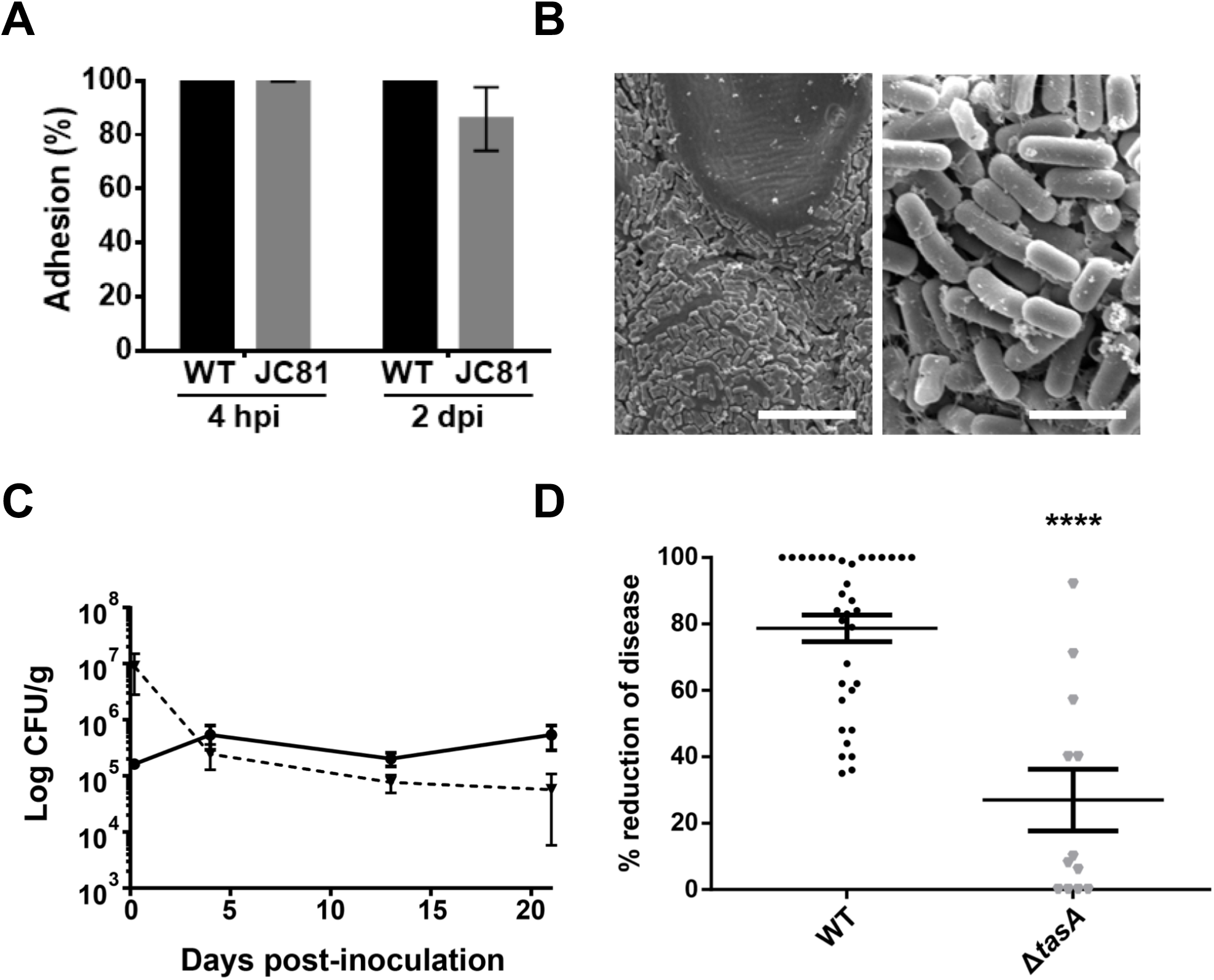
JC81 cells expressing the TasA variant are ecologically compromised on the phylloplane. A) JC81 cells showed similar adhesion to that of the WT cells on melon leaves 4 h or 2 days post-inoculation. Average values of three biological replicates are shown. Error bars represent the SEM. B) Electron micrographs of samples taken 20 days post-inoculation show an intermediate colonization pattern for JC81 compared with those of the WT and *ΔtasA* null mutant strains. C) The persistence of JC81 cells on melon leaves was reduced compared to that of the WT cells. Average values of five biological replicates are shown. Error bars represent the SEM. D) The JC81 strain failed to control the disease induced by the phytopathogenic fungi *P. xanthii*. Average values of three biological replicates are shown. Error bars represent the SEM. Statistical significance was assessed via one-way ANOVA with the Dunnet test (**** p<0.0001).

## Discussion

The ECM provides cells with a myriad of advantages, such as robustness, colony cohesiveness, and protection against external aggression^9, 10, 27^. Studies of *B. subtilis* biofilms have revealed that the ECM is mainly composed of polysaccharides^31^ and the proteins TasA and BslA^27, 32^. TasA is a confirmed functional amyloid that provides structural support to the biofilm in the form of robust fibers^33^. A recent study demonstrated that there is heterogeneity in the secondary structure of TasA; however, in biofilms, its predominant conformation is in the form of stable fibers enriched in β-sheets^34^. In this study, we demonstrate that in addition to its structural role in ECM assembly, TasA is also required for normal colony development – both of which are functions that contribute to the full fitness of *Bacillus* cells on the phylloplane.

The physiological alterations observed in *ΔtasA* null strain reflect a process of progressive cellular deterioration characteristic of senescence^56–58^, including early activation of secondary metabolism, low energy metabolic activity, and accumulation of damaged molecular machinery that is required for vital functions. Indeed, it has been previously demonstrated that such metabolic changes can trigger PCD in other bacterial species, in which over-acidification of the cytoplasm eventually leads to the activation of cell death pathways^54^. Interestingly, cytoplasmic acidification due to the production of acetic acid has been linked to higher ROS generation and accelerated aging in eukaryotes^78^. As mentioned throughout this study, ROS generation leads to ongoing DNA damage accumulation, phospholipid oxidation, and changes in cell membrane potential and functionality, all of which are major physiological changes that eventually lead to declines in cellular fitness and, ultimately, to cell death^79–81^. The fact that we could restore the physiological status of *tasA* null mutant cells by ectopically expressing a mutated TasA protein incapable of rescuing biofilm formation permitted us to separate two roles of TasA: i) its structural function, necessary for ECM assembly; and ii) its cytological functions involved in regulating membrane dynamics and PCD. Our data indicate that this previously unreported function does not involve TasA amyloid fibers or its role in ECM assembly, and that it is more likely related to the TasA found in the DRM of the cell membrane, where the FMMs (like the lipid rafts of eukaryotic cells) are located. It is not unprecedented that amyloid proteins interact with functional domains within the cell membrane. In eukaryotic cells, for instance, it has been reported that lipid rafts participate in the interaction between the amyloid precursor protein and the secretase required for the production of the amyloid-β peptide, which is responsible for Alzheimer’s disease^82^. Indeed, our results are supported by evidence that TasA can preferentially interact with model bacterial membranes, which affects fiber assembly^83^, and that TasA fibers are located and attached to the cell surface via a proposed interaction with TapA, which forms foci that seem to be present on the cell wall^76^. Interestingly, TapA has been recently characterized as a two-domain, partially disordered protein^84^. Disordered domains can be flexible enough to interact with multiple partners^85, 86^, suggesting a similar mechanism for TapA: the N-terminal domain might be involved in the interaction with other protein partners, whereas the C-terminal disordered domain might anchor the protein to the cell surface. All of these observations led us to propose that TasA may drive the stabilization of the FMMs in the cell membrane either directly via interactions with certain phospholipids or indirectly via interactions with other proteins. This model is further supported by the fact that the *ΔtasA* cells show mislocalization of FloT (Fig. 7A), which is typically present in the FMM, and induction of many genes that encode DRM components or other factors that interact with FloT alone or with FloT and FloA (suppl. Table 4).

The physiological alterations observed in the *ΔtasA* strain have ecological implications. The intrinsic stress affecting the mutant cells reduced their ability to survive in natural environments; however, paradoxically, their higher induction of secondary metabolism seemed to indirectly and efficiently target fungal pathogens. This scenario could explain why *ΔtasA* cells, which show clear signs of stress, display efficient biocontrol properties against *P. xanthii*. However, the sharp time-dependent decrease in the *ΔtasA* population on leaves suggests that its antifungal production could be beneficial during short-term interactions, but insufficient to support long-term antagonism unless there is efficient colonization and persistence on the plant surface. We previously speculated that biofilm formation and antifungal production were two complementary tools used by *Bacillus* cells to efficiently combat fungi. Our current study supports this concept, but also enhances our understanding of the roles of the different ECM components. More specifically, we demonstrated that the amyloid protein TasA is the most important bacterial factor during the initial attachment and further colonization of the plant host. Two lines of evidence downplay the importance of the EPS during the early establishment of physical contact. First, an EPS mutant strain behaves similarly to the WT strain, and second, the naturally occurring overexpression of the *eps* genes in the *ΔtasA* were unable to revert the adhesion defect. These observations are more consistent with a role for the EPS, along with BslA, in providing biofilms with protection against external aggressors^31, 87^. A similar role for a functional amyloid protein in bacterial attachment to plant surfaces was found for the *Escherichia coli* curli protein. Transcriptomic studies showed induction of curli expression during the earlier stages of attachment after the cells came into contact with the plant surface, and a curli mutant strain was defective in this interaction^88, 89^. The distinct morphological and biochemical variations typical of amyloids make them perfect candidates for modulating cellular ecology. The observation that *ΔtasA* cells are incapable of colonization in the rhizosphere^90^ clearly indicates the need for more in-depth investigation into these two distinctive ecological niches to understand the true roles of specific bacterial components. In addition to demonstrating enhanced production of antifungal compounds, our study revealed additional features that might contribute to the potency of stressed *Bacillus* cells in arresting fungal growth, in particular the overproduction of acetoin via increased expression of the *alsS* and *alsD* genes. Acetoin is a volatile compound produced via fermentative and overflow metabolism, and it has been demonstrated to mediate communication between beneficial bacteria and plants by activating plant defense mechanisms either locally or over long distances in a phenomenon known as induced systemic resistance (ISR)^91, 92^.

In summary, we have proven that the amyloid protein TasA participates in the proper maturation of *Bacillus* colonies, a function that, along with its previously reported role in ECM assembly, contributes to long-term survival, efficient colonization of the phylloplane, and a competitive advantage against competitors mediated by antifungal production. The absence of TasA leads to a series of physiological changes, likely triggered by alterations in membrane dynamics and effects on the FMMs, including an arrest of cell differentiation^9^ that paradoxically increases the competitiveness of the mutant cells during short-term interactions via their ability to adapt to stress and their cellular response to early maturation. However, lack of TasA reduces cell fitness during mid-to long-term interactions via increased intrinsic cellular stress and the absence of a structured ECM, both of which limit the adaptability of the cells to the stressful phylloplane.

## Material and methods

### Bacterial strains and culture conditions

The bacterial strains used in this study are listed in supplementary table 1. Bacterial cultures were grown at 37 °C from frozen stocks on Luria-Bertani (LB: 1% tryptone (Oxoid), 0.5% yeast extract (Oxoid) and 0.5% NaCl) plates. Isolated bacteria were inoculated in the appropriate medium. The biotrophic fungus *Podosphaera xanthii* was grown at 25 °C from a frozen stock on cucumber cotyledons and maintained on them until inoculum preparation. Biofilm assays were performed on MSgg medium: 100 mM morpholinepropane sulphonic acid (MOPS) (pH 7), 0.5% glycerol, 0.5% glutamate, 5 mM potassium phosphate (pH 7), 50 μg/ml tryptophan, 50 μg/ml phenylalanine, 50 μg/ml threonine, 2 mM MgCl_2_, 700 μM CaCl_2_, 50 μM FeCl_3_, 50 μM MnCl_2_, 2 μM thiamine, 1 μM ZnCl_2_. For the *in vitro* lipopeptide detection and assays with cell-free supernatants, medium optimized for lipopeptide production (MOLP) was used and prepared as previously described^93^. For cloning and plasmid replication, *Escherichia coli* DH5α was used. *Bacillus subtilis* 168 is a domesticated strain used to transform the different constructs into *Bacillus subtilis* NCIB3610. The antibiotic final concentrations for *B. subtilis* were: MLS (1 μg/ml erythromycin, 25 μg/ml lincomycin); spectinomycin (100 μg/ml); tetracycline (10 μg/ml); chloramphenicol (5 μg/ml); kanamycin (10 μg/ml).

### Strain construction

All of the primers used to generate the different strains are listed in Supplementary table 2. To build the strain YNG001, the promoter of the fengycin operon was amplified with the Ppps-ecoRI.F and Ppps-HindIII.R primer pair. The PCR product was digested with EcoRI and HindIII and cloned into the pKM003 vector cut with the same enzymes. The resulting plasmid was transformed by natural competence into *B. subtilis* 168 replacing the *amyE* neutral locus. Transformants were selected via spectinomycin resistance. The same plasmid was used to build the strain YNG002 by transforming a *ΔtasA* strain of *B. subtilis* 168. Strains DR8 and DR9 were constructed similarly by amplifying the promoter of the *epsA-O* operon with the primers Peps-ecoRI.F and Peps-HindIII.R and then cloning the insert into pKM003. The plasmid was transformed into the WT (DR8) and *ΔtasA* (DR9) *B. subtilis* 168 strains.

Strain YNG003 was constructed using the primers amyEUP-Fw, amyEUP-Rv, Ppps-Fw, Ppps-Rv, Yfp-Fw, Yfp-Rv, Cat-Fw. Cat-Rv, amyEDOWN-Fw, amyEDOWN-Rv to separately amplify the relevant fragments. The fragments were then joined using the NEB builder HiFi DNA Assembly Master Mix (New England Biolabs). The construct was made using pUC19 digested with BamHI as the vector backbone. The final plasmid was then transformed into *B. subtilis* 168 replacing *amyE,* and transformants were selected via chloramphenicol resistance.

Strain JC97 was generated using the primers bslAUP-Fw, bslADOWN-Rv, Spc-Fw, Spc-Rv, bslaUP-Fw and bslADOWN-Rv and XbaI-digested pUC19 as the vector backbone. The fragments were assembled using NEB Builder HiFi DNA Assembly Master Mix.

Strains JC70, JC81 and JC149 were constructed via site-directed mutagenesis (QuickChange Lightning Site Directed Mutagenesis Kit – Agilent Technologies). Briefly, the *tapA* operon (*tapA-sipW-tasA*), including its promoter, was amplified using the primers TasA_1_mutb and YSRI_2, and the resulting product was digested with BamHI and SalI and cloned into the pDR183 vector^94^. Next, the corresponding primers (see suppl. Table 5A) were used to introduce the alanine substitution mutations into the desired positions of the TasA amino acid sequence. The entire plasmid was amplified from the position of the primers using Pfu DNA polymerase. The native plasmid, which was methylated and lacked the mutations, was digested with DpnI enzyme. The plasmids containing the native version of TasA (JC70) or the mutated versions (JC81 and JC149) were transformed into the *B. subtilis* 168 *Δ(tapA-sipW-tasA)* strain replacing the *lacA* neutral locus. Genetic complementation was observed in strain JC70 as a control. Transformants were selected via MLS resistance.

All of the *B. subtilis* strains generated were constructed by transforming *B. subtilis* 168 via its natural competence and then using the positive clones as donors for transferring the constructs into *Bacillus subtilis* NCIB3610 via generalized SPP1 phage transduction^95^.

### Biofilm assays

To analyze colony morphology under biofilm-inducing conditions, we used a method described elsewhere^96^. Briefly, the bacterial strains were grown on LB plates overnight at 37 °C, and the resulting colonies were resuspended in sterile distilled water at an OD_600_ of 1. Next, 2-µl drops of the different bacterial suspensions were spotted on MSgg agar plates and incubated at 30 °C. Colonies were removed at the appropriate time points (24, 48 and 72 h) for the different analyses.

For the CFU counts of the colonies from the different strains, 24-, 48- and 72-hour-old colonies grown on MSgg agar plates were removed, resuspended in 1 ml of sterile distilled water, and subjected to mild sonication (three rounds of 20 second pulses at 20% amplitude). The resulting suspensions were serially diluted and plated to calculate the CFUs per colony (total CFU). To estimate the CFUs corresponding to sporulated cells (CFU endospores), the same dilutions were heated at 80 °C for 10 minutes and plated. The sporulation percentage was calculated as (CFU endospores/total CFU)*100.

### Biofilm fractionation

To analyze the presence of TasA in the different strains, biofilms were fractionated into cells and ECM as described elsewhere^96^, and both fractions were analyzed separately. Briefly, 72-hour-old colonies grown under biofilm-inducing conditions on MSgg-agar plates were carefully lifted from the plates and resuspended in 10 ml of MS medium (MSgg broth without glycerol and glutamate, which were replaced by water) with a 25 ^5/8^ G needle. Next, the samples were subjected to mild sonication in a Branson 450 digital sonifier (4-5 5 seconds pulses at 20% amplitude) to ensure bacterial resuspension. The bacterial suspensions were centrifuged at 9000 g for 20 minutes to separate the cells from the extracellular matrix. The cell fraction was resuspended in 10 ml of MS medium and stored at 4 °C until further processing. The ECM fraction was filtered through a 0.22 µm filter and stored at 4 °C.

For protein precipitation, 2 ml of the cell or ECM fractions were used. The cell fraction was treated with 0.1 mg/ml lysozyme for 30 minutes at 37 °C. Next, both fractions were treated with a 10% final concentration of trichloroacetic acid and incubated in ice for 1 h. Proteins were collected by centrifugation at 13,000 g for 20 minutes, washed twice with ice-cold acetone, and dried in an Eppendorf Concentrator Plus 5305 (Eppendorf).

### Cell membrane fractionation

Crude membrane extracts were purified from 50 ml MSgg liquid cultures (with shaking) of the different *B. subtilis* strains. Cultures were centrifuged at 7,000 g for 10 minutes at 4 °C and then resuspended in 10 ml of PBS. Lysozyme was added at a final concentration of 20 µg/ml and the cell suspensions were incubated at 37 °C for 30 minutes. After incubation, the lysates were sonicated on ice with a Branson 450 digital sonifier using a cell disruptor tip and 45 second pulses at 50% amplitude with pauses of 30 seconds between pulses until the lysates were clear. Next, the cell lysates were centrifuged at 10,000 g for 15 minutes to eliminate cell debris, and the supernatants were separated and passed through a 0.45 µm filter. To isolate the cell membrane, the filtered lysate was ultracentrifuged at 100,000 g for 1 hour at 4 °C. The supernatant, which contained the cytosolic proteins, was separated and kept at −20 °C. The pellet, which contained the crude membrane extract, was washed 3 times with PBS and processed using the CelLytic MEM protein extraction kit from Sigma. Briefly, the membrane fractions were resuspended in 600 µl of lysis and separation working solution (lysis and separation buffer + protease inhibitor cocktail) until a homogeneous suspension was achieved. Next, the suspension was incubated overnight at 4 °C on a stirring wheel. After incubation, the suspension is incubated at 37 °C for 30 minutes and then centrifuged at 3,000 g for 3 minutes. The DSM (upper phase) was separated and kept at −20 °C, and the DRM (lower phase) was washed three times with 400 µl of wash buffer by repeating the process from the 37 °C incubation step. Three washes were performed to ensure the removal of all hydrophilic proteins. The isolated DRM was kept at −20 °C until use. The DRM, DSM and cytosolic fractions were used directly for immunodetection.

### SDS-PAGE and immunodetection

Precipitated proteins were resuspended in 1x Laemmli sample buffer (BioRad) and heated at 100 °C for 5 minutes. Proteins were separated via SDS-PAGE in 13% acrylamide gels and then transferred onto PVDF membranes using the Trans-Blot Turbo Transfer System (BioRad) and PVDF transfer packs (BioRad). For immunodetection of TasA, the membranes were probed with anti-TasA antibody (rabbit) used at a 1:20,000 dilution in Pierce Protein-Free (TBS) blocking buffer (ThermoFisher). For immunodtection of FloT-YFP, a commercial anti-GFP primary antibody (Clontech living colors full-length polyclonal antibody) developed in rabbit was used at a 1:1,000 dilution in the buffer mentioned above. A secondary anti-rabbit IgG antibody conjugated to horseradish peroxidase (BioRad) was used at a 1:3,000 dilution in the same buffer. The membranes were developed using the Pierce ECL Western Blotting Substrate (ThermoFisher).

### Bioassays on melon leaves

Bacterial strains were grown in liquid LB at 30 °C overnight. The cells in the cultures were washed twice with sterile distilled water. The bacterial cell suspensions were adjusted to the same OD_600_ and sprayed onto leaves of 4- to 5-week-old melon plants. Two hours later, a suspension of *P. xanthii* conidia was sprayed onto each leaf at a concentration of 4−10×10^4^ spores/ml. The plants were placed in a greenhouse or in a growth chamber at 25 °C with a 16 h photoperiod, 3800 lux and 85% RH. The severity of the symptoms was evaluated as the percentage of leaf covered by powdery mildew, as previously described^97^. Briefly, the entire leaf area and the powdery mildew damage area were measured using FiJi^98^, and the ratio of infection was calculated using the formula [(damage area/leaf area)*100].

The persistence of bacterial strains on plant leaves was calculated via CFU counts performed over the twenty-one days following inoculation. Three different leaves from three different plants were individually placed into sterile plastic stomacher bags and homogenized in a lab blender (Colworth Stomacher-400, Seward, London, UK) for 3 min in 10 ml of sterile distilled water. The leaf extracts were serially diluted and plated to calculate the CFUs at each time point. The plates were incubated at 37 °C for 24 h before counting.

The adhesion of bacterial cells to melon leaves was estimated by comparing the number of cells released from the leaf versus the cells attached to the surface. The surfaces of individual leaves were placed in contact with 100 ml of sterile distilled water in glass beakers and, after 10 minutes of stirring (300 rpm), the water and leaf were plated separately. The leaves were processed as described above. Adhesion was calculated as the ratio: (water CFU/total CFU)*100. The data from all of the different strains were normalized to the result of the WT strain (100% adhesion).

### Antifungal activity of cell-free supernatant against Podosphaera xanthii

*B. subtilis* strains were grown for 72 h at 30 °C in medium optimized to encourage lipopeptide production (MOLP), and the supernatant was centrifuged and filtered (0.22 µm). One-week-old cotyledons were disinfected with 20% commercial bleach for 30 seconds and then submerged two times in sterile distilled water for 2 minutes and then air dried. 10-mm discs were excised with a sterilized cork borer, incubated with cell-free supernatants for 2 h, and then left to dry. Finally, the discs were inoculated with *P. xanthii* conidia as previously described^99^.

### Lipopeptides production analysis

For the *in vitro* lipopeptide detection, bacteria were grown in MOLP for 72 h. The cultures were centrifuged, and the supernatants were filtered (0.22 µm) prior to analysis via MALDI-TOF/TOF. For *in situ* lipopeptide detection on inoculated leaves, leaf discs were taken 21 days post-inoculation with a sterile cork borer and then placed directly on an UltrafleXtreme MALDI plate. A matrix consisting of a combination of CHCA (α-cyano-4-hydroxycinnamic acid) and DHB (2,5-dihydroxybenzoic acid) was deposited over the discs or the supernatants (for the *in vitro* culture), and the plates were inserted into an UltrafleXtreme MALDI-TOF/TOF mass spectrometer. The mass spectra were acquired using the Bruker Daltonics FlexControl software and were processed using Bruker Daltonics FlexAnalysis.

### Electron microscopy analysis

For the scanning electron microscopy analysis, leaf discs were taken 21 days post-inoculation as previously described and fixed in 0.1 M sodium cacodylate and 2% glutaraldehyde overnight at 4 °C. Three washes were performed with 0.1 M sodium cacodylate and 0.1 M sucrose followed by ethanol dehydration in a series of ethanol solutions from 50% to 100%. A final drying with hexamethyldisilazane was performed as indicated^100^. The dried samples were coated with a thin layer of iridium using an Emitech K575x turbo sputtering coater before viewing in a Helios Nanolab 650 Scanning Electron Microscope and Focus Ion Beam (SEM-FIB) with a Schottky-type field emission electron gun.

For the transmission electron microscopy analysis, bacterial colonies grown on MSgg agar for the appropriate times were fixed directly using a 2% paraformaldehyde-2.5% glutaraldehyde-0.2 M sucrose mix in phosphate buffer 0.1 M (PB) overnight at 4 °C. After three washes in PB, portions were excised from each colony and then post-fixed with 1% osmium tetroxide solution in PB for 90 minutes at room temperature, followed by PB washes, and 15 minutes of stepwise dehydration in an ethanol series (30%, 50%, 70%, 90%, and 100% twice). Between the 50% and 70% steps, colonies were incubated *in-bloc* in 2% uranyl acetate solution in 50% ethanol at 4 °C, overnight. Following dehydration, the samples were gradually embedded in low-viscosity Spurr’s resin: resin:ethanol, 1:1, 4 hours; resin:ethanol, 3:1, 4 hours; and pure resin, overnight. The sample blocks were embedded in capsule molds containing pure resin for 72 h at 70 °C. The samples were visualized under a FEI Tecnai G^2^ 20 TWIN Transmission Electron Microscope at an accelerating voltage of 80 KV. The images were taken using a side-mounted CCD Olympus Veleta with 2k x 2k Mp.

### Whole-transcriptome analysis and qRT-PCR

Biofilms were grown on MSgg agar as described above. After 72 h of growth, colonies were recovered and stored at −80 °C. All of the assays were performed in duplicate. The collected cells were resuspended and homogenized via passage through a 25 ^5/8^ G needle in BirnBoim A^101^ buffer (20% sucrose, 10 mM Tris-HCl pH 8, 10 mM EDTA and 50 mM NaCl). Lysozyme (10 mg/ml) was added, and the mixture was incubated for 30 minutes at 37 °C. After disruption, the suspensions were centrifuged, and the pellets were resuspended in Trizol reagent (Invitrogen). Total RNA extraction was performed as instructed by the manufacturer. DNA removal was carried out via in-column treatment with the rDNAse included in the Nucleo-Spin RNA Plant Kit (Macherey-Nagel) following the instructions of the manufacturer. The integrity and quality of the total RNA was assessed with an Agilent 2100 Bioanalyzer (Agilent Technologies) and by gel electrophoresis.

To perform the RNA sequencing analysis, rRNA removal was performed using the RiboZero rRNA removal (bacteria) Kit from Illumina, and 100-bp single-end read libraries were prepared using the TruSeq Stranded Total RNA Kit (Illumina). The libraries were sequenced using a NextSeq550 instrument (Illumina). The data analysis was performed using fastQC for sample quality control, EdgePRO^102^ for mapping against reference genomes (*B. subtilis 168* was used for NCIB3610, Genbank: NC_000964.3) and quantifying gene expression, NOIseq^103^ to normalize the samples, and edgeR^102^ for differential expression analysis. Genes were considered differentially expressed when their logFC was higher than 1 or lower than −1 with a p-value < 0.05. The data were deposited in the GEO database (GEO accession GSE124307).

Quantitative real-time (qRT)-PCR was performed using the iCycler-iQ system and the iQ SYBR Green Supermix Kit from Bio-Rad. The primer pairs used to amplify the target genes were designed using the Primer3 software (http://bioinfo.ut.ee/primer3/) and Beacon designer (http://www.premierbiosoft.com/qOligo/Oligo.jsp?PID=1), maintaining the parameters described elsewhere^104^. For the qRT-PCR assays, the RNA concentration was adjusted to 100 ng/µl. Next, 1 µg of DNA-free total RNA was retro-transcribed into cDNA using the SuperScript III reverse transcriptase (Invitrogen) and random hexamers in a final reaction volume of 20 µl according to the instructions provided by the manufacturer. The qRT-PCR cycle was: 95 °C for 3 min, followed by PCR amplification using a 40-cycle amplification program (95 °C for 20 sec, 56 °C for 30 sec, and 72 °C for 30 sec), followed by a third step of 95 °C for 30 sec. To evaluate the melting curve, 40 additional cycles of 15 sec each starting at 75 °C with stepwise temperature increases of 0.5 °C per cycle were performed. To normalize the data, the *rpsJ* gene, encoding the 30S ribosomal protein S10, was used as a reference gene^105^. The target genes *fenD*, encoding fengycin synthetase D, *alsS*, encoding acetolactate synthase, *albE*, encoding bacteriocin subtilosin biosynthesis protein AlbE, *bacB*, encoding the bacilysin biosynthesis protein BacB, and *srfAA* encoding surfactin synthetase A, were amplified using the primer pairs given in supplementary Table 3, resulting in the generation of fragments of 147 bp, 82 bp, 185 bp, 160 bp and 9 4bp, respectively. The primer efficiency tests and confirmation of the specificity of the amplification reactions were performed as previously described^106^. The relative transcript abundance was estimated using the *ΔΔ* cyclethreshold (Ct) method^107^. Transcriptional data of the target genes was normalized to the *rpsJ* gene and shown as the fold-changes in the expression levels of the target genes in each *B. subtilis* mutant strain compared to those in the WT strain. The relative expression ratios were calculated as the difference between the qPCR threshold cycles (Ct) of the target gene and the Ct of the *rpsJ* gene (*Δ*Ct= Ct*rgene of interest* – Ct*rpsJ*). Fold-change values were calculated as 2^-*ΔΔ*Ct^, assuming that one PCR cycle represents a two-fold difference in template abundance^108, 109^. The qRT-PCR analyses were performed three times (technical replicates) using three independent RNA isolations (biological replicates).

### Flow cytometry assays

Cells were grown on MSgg agar at 30 °C. At different time points, colonies were recovered in 500 μL of PBS and resuspended with a 25 ^5/8^ G needle. For the promoter expression assays, colonies were gently sonicated as described above to ensure complete resuspension, and the cells were fixed in 4% paraformaldehyde in PBS and washed three times in PBS. To evaluate the physiological status of the different *B. subtilis* strains, cells were stained for 30 minutes with 5 mM 5-cyano-2,3-ditolyltetrazolium chloride (CTC) and 15 µM 3-(p-hydroxyphenyl) fluorescein (HPF).

The flow cytometry runs were performed with 200 μl of cell suspensions in 800 μL of GTE buffer (50 mM glucose, 10 mM EDTA, 20 mM Tris-HCl; pH 8), and the cells were measured on a Beckman Coulter Gallios™ flow cytometer using 488 nm excitation. YFP and HPF fluorescence were detected with 550 SP or 525/40 BP filters. CTC fluorescence was detected with 730 SP and 695/30BP filters. The data were collected using Gallios™ Software v1.2 and further analyzed using Kaluza Analysis v1.3.

### Intracellular pH analysis

Intracellular pH was measured as previously described^54^. Briefly, colonies of the different strains grown on MSgg agar at 30 °C were taken at different time points and recovered in potassium phosphate buffer (PPB) pH 7 and gently sonicated as described above. Next, the cells were incubated in 10 µl of 1 mM 5-(6)carboxyfluorescein diacetate succinimidyl (CFDA) for 15 minutes at 30 °C. PPB supplemented with glucose (10 mM) was added to the cells for 15 minutes at 30 °C to remove the excess dye. After two washes with the same buffer, the cells were resuspended in 50 mM PPB (pH 4.5).

Fluorescence was measured in a FLUOstar Omega (BMG labtech) microplate spectrofluorometer using 490nm/525nm as the excitation and emission wavelengths, respectively. Conversion from the fluorescence arbitrary units into pH units was performed using a standard calibration curve.

### Fluorescence microscopy

The localization of TasA in *B. subtilis* protoplasts was evaluated using a TasA-mCherry translational fusion (see suppl. Table 5). To generate the protoplast cells, *B. subtilis* colonies of the different strains grown on MSgg agar plates for 72 h were resuspended in protoplast buffer (20 mM potassium phosphate, pH 7.5, 15 mM MgCl_2_, 20% sucrose), mildly sonicated as describe above and incubated for 30 min in the presence of 10 µg/ml lysozyme at 37 °C. The protoplast suspensions were mounted and visualized immediately under a Leica DM2500 LED fluorescence microscope with standard Texas Red (TX2 Ex. 560/40 Em. 645/76) filter to visualize cells expressing the *tasA-mCherry* construct. The images were taken with a Leica DFC 7000T 2.8 MP camera.

FM 4-64 was purchased from ThermoFischer and was used at a final concentration of 5 µg/ml and was visualized using a standard GFP filter (GFP Ex. 470/40 Em. 525/50)

### Confocal laser scanning microscopy (CLSM)

Cell death in the bacterial colonies was evaluated using the LIVE/DEAD BacLight Bacterial Viability Kit (Invitrogen). Equal volumes of both components included in the kit were mixed, and 2 µl of this solution was used to stain 1 ml of the corresponding bacterial suspension. Sequential acquisitions were configured to visualize the live or dead bacteria in the samples. Acquisitions with excitation at 488 nm and emission recorded from 499 to 554 nm were used to capture the images from live bacteria, followed by a second acquisition with excitation at 561 nm and emission recorded from 592 to 688 nm for dead bacteria.

For the microscopic analysis and quantification of lipid peroxidation in live bacterial samples, we used the image-iT Lipid Peroxidation Kit (Invitrogen) following the manufacturer’s instructions with some slight modifications. Briefly, colonies of the different strains were grown on MSgg plates at 30 °C, isolated at different time points, and resuspended in 1 ml of liquid MSgg medium as described in the previous sections. 5 mM cumene hydroperoxide (CuHpx)-treated cell suspensions of the different strains at the corresponding times were used as controls. The cell suspensions were then incubated at 30 °C for 2 h and then stained with a 10 µM solution of the imageIT lipid peroxidation sensor for 30 minutes. Finally, the cells were washed three times with PBS, mounted, and visualized immediately. Images of the stained bacteria were acquired sequentially to obtain images from the oxidized to the reduced states of the dye. The first image (oxidized channel) was acquired by exciting the sensor at 488 nm and recording the emissions from 509 to 561 nm, followed by a second acquisition (reduced channel) with excitement at 561 nm and recording of the emissions from 590 to 613 nm.

Membrane potential was evaluated using the image-iT TMRM (tetramethylrhodamine, methyl ester) reagent (Invitrogen) following the manufacturer’s instructions. Colonies grown at 30 °C on MSgg solid medium were isolated at different time points and resuspended as described above. Samples treated prior to staining with 20 µM carbonyl cyanide m-chlorophenyl hydrazine (CCCP), a known protonophore and uncoupler of bacterial oxidative phosphorylation, were used as controls (Fig. S11). The TMRM reagent was added to the bacterial suspensions to a final concentration of 100 nM, and the mixtures were incubated at 37 °C for 30 minutes. After incubation, the cells were immediately visualized by CLSM with excitation at 561 nm and emission detection between 576 and 683 nm.

The amounts of DNA damage in the *B. subtilis* strains at the different time points were evaluated via terminal deoxynucleotidyl transferase (TdT) dUTP Nick-End Labeling (TUNEL) using the *In-Situ* Cell Death Detection Kit with fluorescein (Roche) according to the manufacturer’s instructions. *B. subtilis* colonies were resuspended in PBS and processed as described above. The cells were centrifuged and resuspended in 1% paraformaldehyde in PBS and fixed at room temperature for one hour on a rolling shaker. The cells were then washed twice in PBS and permeabilized in 0.1% Triton X-100 and 0.1% sodium citrate for 30 minutes at room temperature with shaking. After permeabilization, the cells were washed twice with PBS and the pellets were resuspended in 50 µl of the TUNEL reaction mixture (45 µl label solution + 5 µl enzyme solution), and the reactions were incubated for one hour at 37°C in the dark with shaking. Finally, the cells were washed twice in PBS, counterstained with DAPI (final concentration 500 nM), mounted, and visualized by CLSM with excitation at 488 nm and emission detection between 497 and 584 nm.

Membrane fluidity was evaluated via Laurdan generalized polarization (GP) as described previously^110^ with some modifications. Colonies of the different *B. subtilis* strains were grown and processed as described above. The colonies were resuspended in 50 mM Tris pH 7.4 with 0.5% NaCl. Laurdan reagent (6-dodecanoyl-N,N-dimethyl-2-naphthylamine) was purchased from Sigma-Aldrich (Merck) and dissolved in N,N-dimethylformamide (DMF). Samples treated prior to staining with 2% benzyl alcohol, a substance known to increase lipid fluidity^111^, were used as positive controls (Fig. S14). Laurdan was added to the bacterial suspensions to a final concentration of 100 µM. The cells were incubated at room temperature for 10 minutes, mounted, and then visualized immediately using two-photon excitation with a Spectraphysics MaiTai Pulsed Laser tuned to 720 nm (roughly equivalent to 360 nm single photon excitation), attached to a Leica SP5 microscope. Emissions between 432 and 482 nm (gel phase) and between 509 to 547 nm (liquid phase) were recorded using the internal PMT detectors.

The localization of FloT in *B. subtilis* cells was evaluated using a FloT-YFP translational fusion (see suppl. Table 5). Colonies grown at 30 °C on MSgg solid medium were isolated at different time points and resuspended as described above. Samples were mounted and visualized immediately with excitation at 514 nm and emission recorded from 518 to 596 nm.

All images were obtained by visualizing the samples using an inverted Leica SP5 system with a 63x NA 1.4 HCX PL APO oil-immersion objective. For each experiment, the laser settings, scan speed, PMT or HyD detector gain, and pinhole aperture were kept constant for all of the acquired images.

### Image analysis

Image processing was performed using Leica LAS AF (LCS Lite, Leica Microsystems) and FIJI/ImageJ software.

Images of live and dead bacteria from viability experiments were processed automatically, counting the number of live (green) or dead (red) bacteria in their corresponding channels. The percentage of dead cells was calculated dividing the number of dead cells by the total number of bacteria found on a field.

For processing the lipid peroxidation images, images corresponding to the reduced and oxidized channels were smoothed and a value of 3 was then subtracted from the two channels to eliminate the background. The ratio image was calculated by dividing the processed reduced channel by the oxidized channel using the FiJi image calculator tool. The ratio images were pseudo-colored using a color intensity look-up table (LUT), and intensity values of min. 0 and max. 50 were selected. All of the images were batch processed with a custom imageJ macro, in which the same processing options were applied to all of the acquired images. Quantification of the lipid peroxidation was performed in Imaris v7.4 (Bitplane) by quantifying the pixel intensity of the ratio images with the Imaris “spots” tool.

The Laurdan GP acquisitions were processed similarly. Images corresponding to the gel phase channel and the liquid phase channel were smoothed and a value of 10 was subtracted to eliminate the background. The Laurdan GP image was then calculated by applying the following formula:

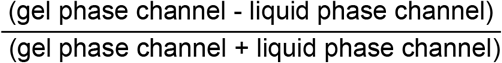

The calculation was performed step by step using the FiJi image calculator tool. Pixels with high Laurdan GP values, typically caused by residual background noise, were eliminated with the “Remove outliers” option using a radius of 4 and a threshold of 5. Finally, the Laurdan GP images were pseudo-colored using a color intensity LUT, and intensity values of min. 0 and max. 1.5 were selected. This processing was applied to all of the acquisitions for this experiment. To quantify the Laurdan GP, bright field images were used for thresholding and counting to create counts masks that were applied to the Laurdan GP images to measure the mean Laurdan GP value for each bacterium.

TUNEL images were analyzed by subtracting a value of 10 in the TUNEL channel to eliminate the background. The DAPI channel was then used for thresholding and counting as described above to quantify the TUNEL signal. The same parameters were used to batch process and quantify all of the images.

To quantify the membrane potential, the TMRM assay images were analyzed as described above using the bright field channel of each image for thresholding and counting to calculate the mean fluorescence intensity in each bacterium. Endospores, which exhibited a bright fluorescent signal upon TMRM staining, were excluded from the analysis. This processing was applied to all of the acquisitions for this experiment.

To quantify the fluorescence of the bacteria expressing the *floT-yfp* construct, images were analyzed as described above using the bright field channel of each image for thresholding and counting to calculate the mean fluorescence intensity in each bacterium.

### Statistical analysis

All of the data are representative of at least three independent experiments with at least three technical replicates. The results are expressed as the mean ± standard error of the mean (SEM). Statistical significance was assessed by performing the appropriate tests (see the figure legends). All analyses were performed using GraphPad Prism version 6. P-values <0.05 were considered significant. Asterisks indicate the level of statistical significance: **\*** = p <0.05, **\*\*** = p <0.01, **\*\*\*** = p <0.001, and **\*\*\*\*** = p<0.0001.

## Acknowledgments

We thank the Ultrasequencing Unit of the SCBI-UMA for RNA sequencing, Juan Felix López Tellez at Bionand for his technical support in the electron transmission analysis, and the flow cytometry service at Bionand. C.M.S is funded by the program Juan de la Cierva Formación (FJCI-2015-23810). This work was supported by grants from ERC Starting Grant (BacBio 637971) and Plan Nacional de I+D+I of Ministerio de Economía y Competitividad (AGL2016-78662-R).

## Author contributions

D.R. conceived the study.

D.R., and J.C.A. and Y.N. designed the experiments.

J.C.A. and Y.N. performed the main experimental work.

M.C.P.B. gave support to some physiological experiments and did q-RT-PCR experiments.

C.M.S. and L.D.M. analyzed and processed the whole transcriptomes.

J.C.A. and J.P. performed and designed the confocal microscopy work and data analysis.

D.R., J.C.A. and Y.N. wrote the manuscript.

D.R., J.C.A., C.M.S., A.V, A.P.G. and L.D.M. contributed critically to writing the final version of the manuscript.

## Competing interests

The authors declare no competing interests

