## Supplementary material for "Dual functionality of the TasA amyloid protein in *Bacillus* physiology and fitness on the phylloplane"

Fig. Sup. 1

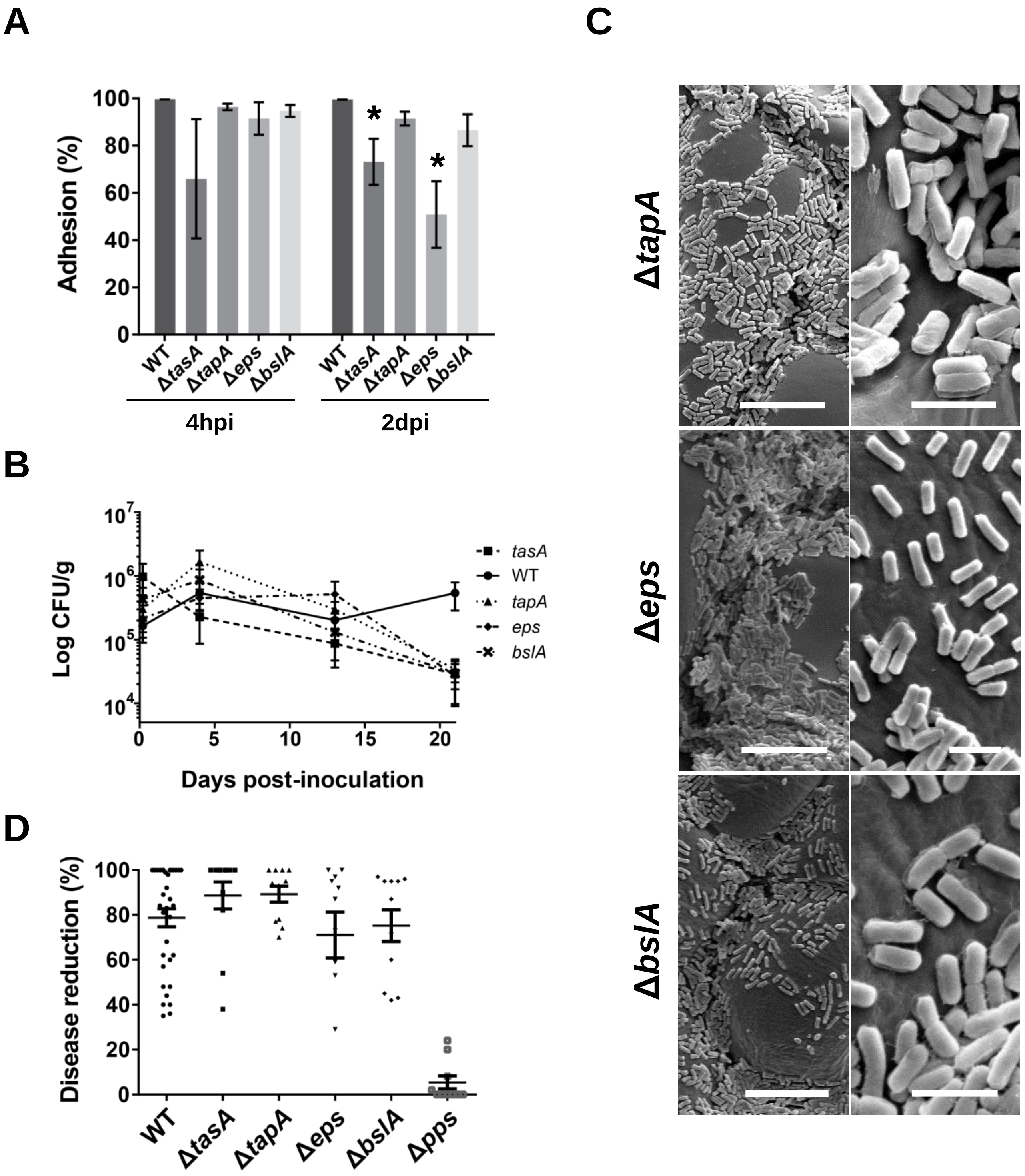


**Figure S1**. **Interaction of *Bacillus subtilis* strains with single mutations in ECM components with melon leaves.** A) Adhesion of the ECM single mutants to melon leaves 4 h and 2 days post-inoculation (hpi and dpi respectively). The *tasA* and *eps* mutants were the only matrix single mutant strains whose adhesion to melon leaves was significantly affected 2 days post-inoculation (dpi). Average values of three biological replicates are shown. Error bars represent the SEM. Statistical significance was assessed via one-way ANOVA with the Dunnet test (* $p<0.05$). B) The *Δeps*, *ΔbslA* and *ΔtapA* strains showed a colonization pattern in which the cells were distributed randomly over the leaf surface with no connection between the cells or extracellular material. Scale bars = 12.5 µm (left) or 2.5 µm (right). C) All of the matrix single mutants showed decreased persistence on plant leaves D) Strains with *tasA* or *tapA* mutations (i.e., no amyloid fibers) showed as much biocontrol activity as that of the WT strain. The *eps* and *bslA* mutant strains showed reduced antagonistic activity compared to that of the WT strain. Average values of three biological replicates are shown. Error bars represent the SEM.

Fig. Sup. 2

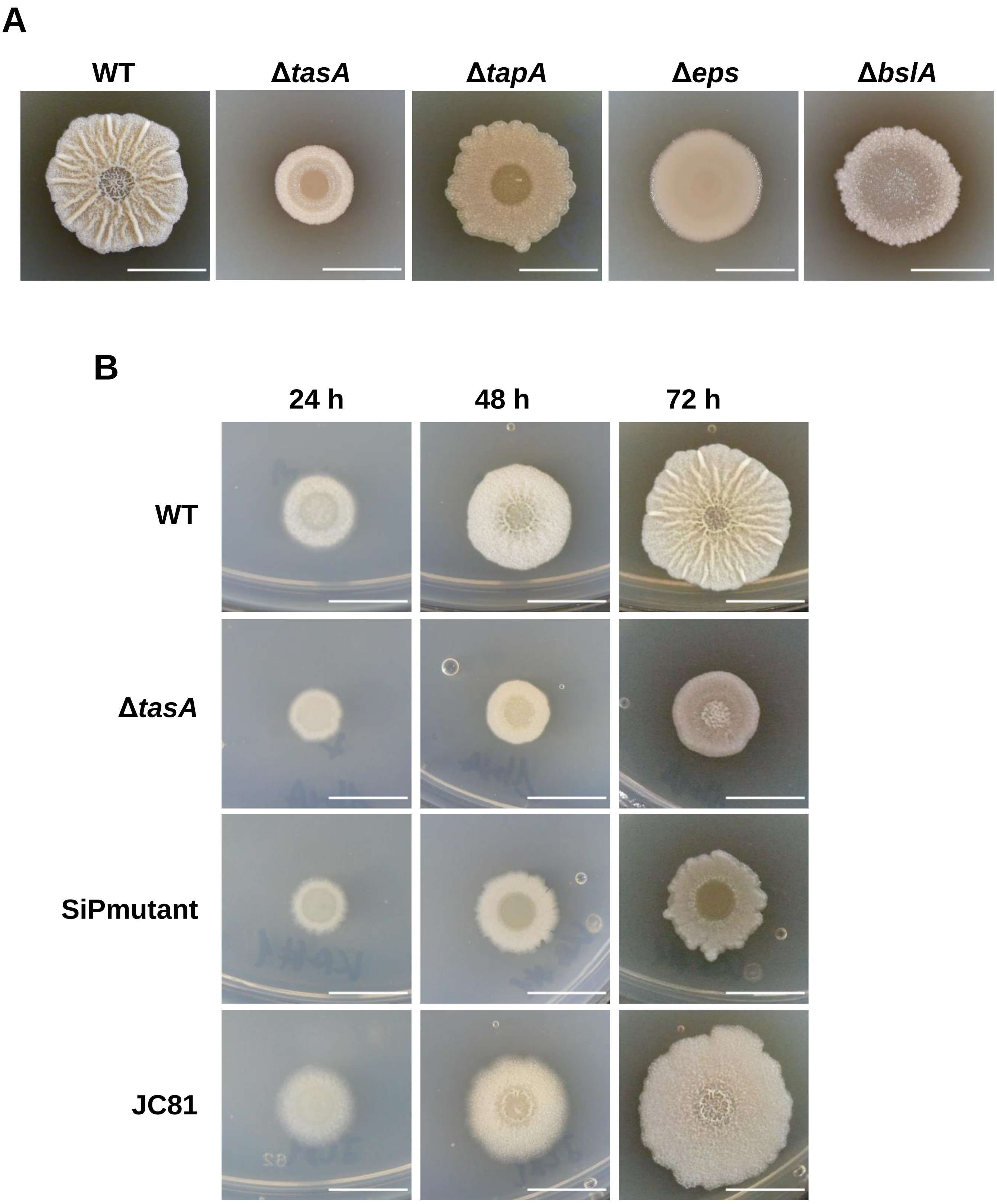


**Figure S2. Morphology of the *Bacillus subtilis* matrix mutants grown under biofilm-inducing conditions**. A) Colony morphologies of the ECM single mutant strains after 72 h of growth on MSgg agar plates. B) Time course of the colony morphologies of the WT, *ΔtasA*, SiPmutant and JC81 strains. Scale bars = 1 cm

**Fig. Sup. 3**

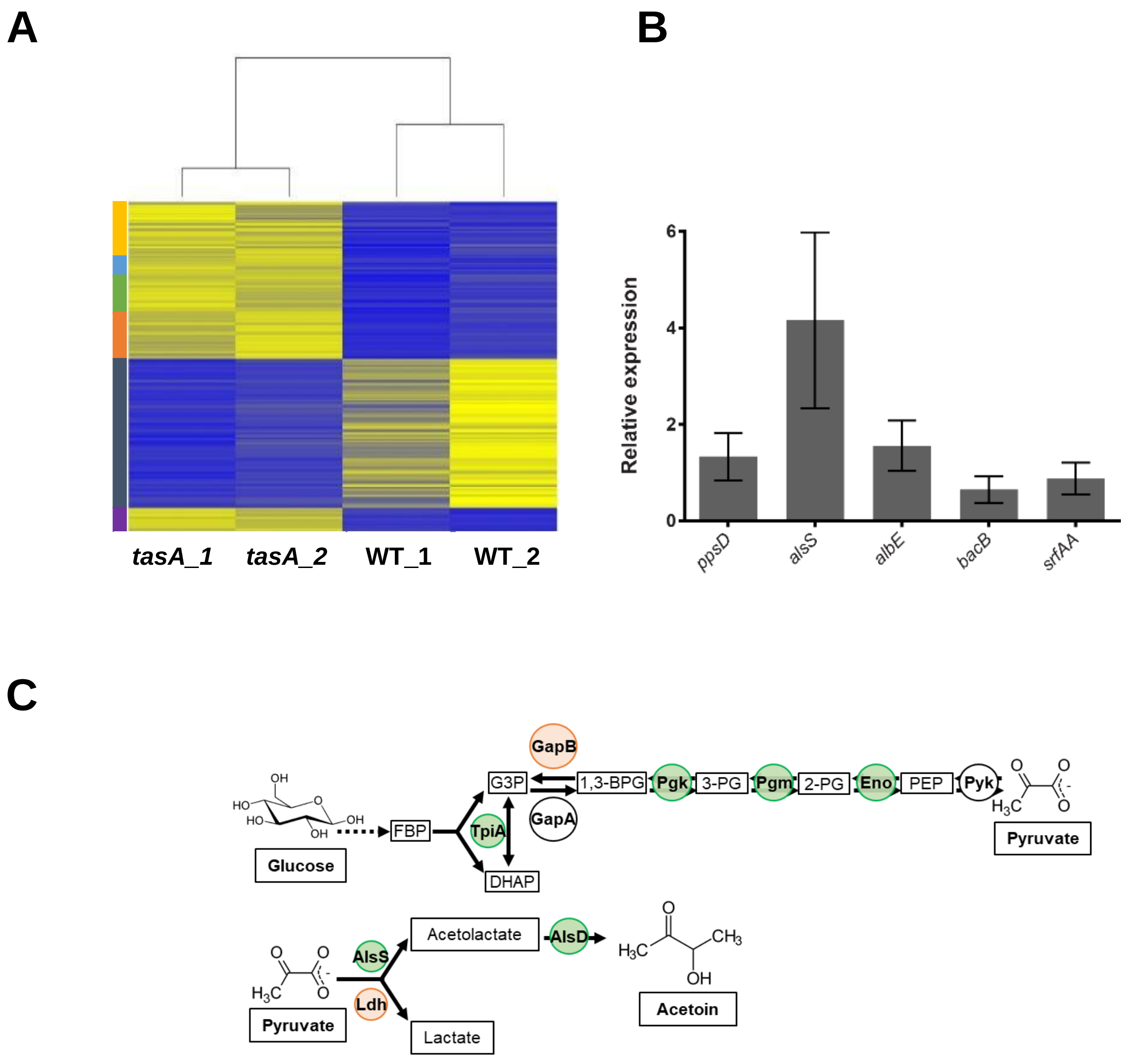


**Figure S3. Deletion of *tasA* had pleiotropic effects on gene expression.** A) Heatmap showing the expression levels of some genes grouped in different categories. Blue indicates repression, and yellow indicates induction. The color bar on the left indicates the different categories: yellow = production of secondary metabolites, cyan = general stress response, green = metabolism of modified versions of cysteine, orange = bacteriophage PBSX, dark blue = sporulation, and purple = biofilm formation. B) qRT-PCR of the expression levels of the *ppsD, alsS, albE, bacB* and *srfAA* genes showed their induction in the Δ*tasA* strain and confirmed the results obtained in the RNA-seq analysis. C-E) The diagram shows the pathway that terminates in acetoin synthesis. Some steps of the glycolysis pathway are induced in the Δ*tasA* strain, whereas the divergent or gluconeogenetic steps are repressed.

### Fig. Sup. 4

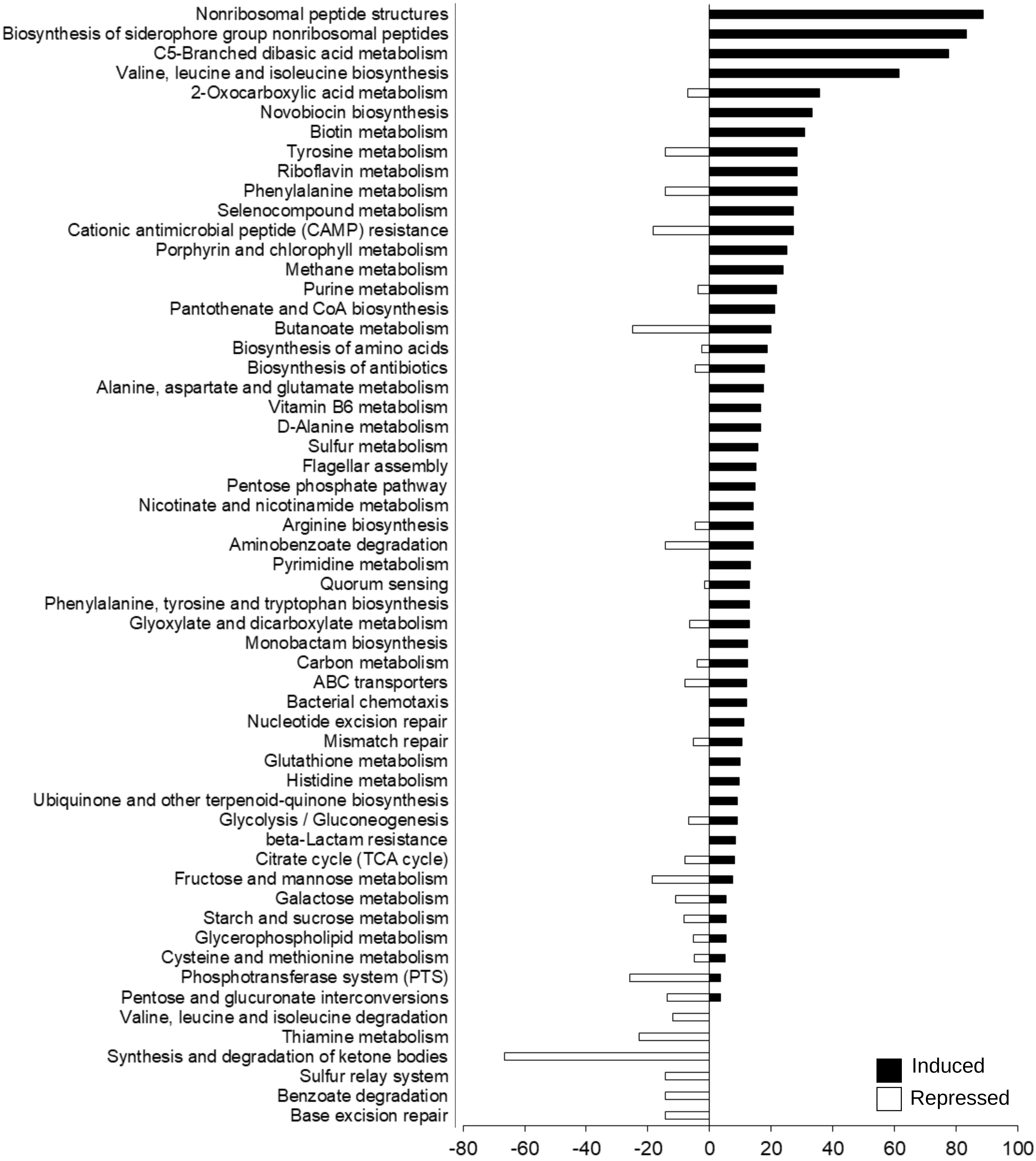


**Figure S4. Multiple pathways were altered in *ΔtasA* cells.** KEGG pathway analysis of the genes that were induced (black bars) or repressed (white bars) in the *ΔtasA* strain compared to the WT strain after 72 h of growth on MSgg agar plates. The analysis was performed with KOBAS 3.0.

**Fig. Sup. 5**

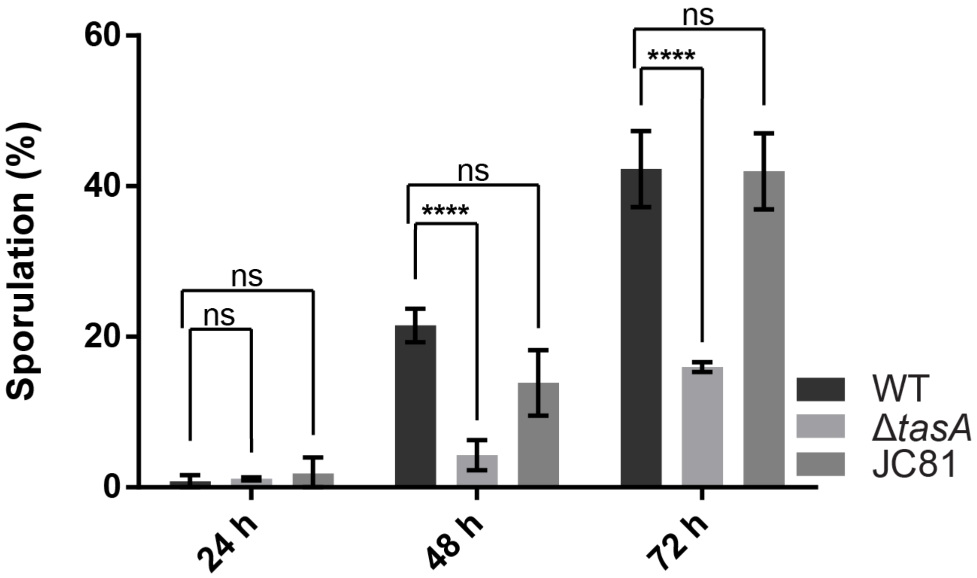


**Figure S5. The *ΔtasA* strain shows a lower percentage of sporulation compared to those in the WT and JC81 strains.** Sporulation rates of the WT, *ΔtasA* and JC81 colonies 24, 48 and 72 h. The *ΔtasA* colonies showed a sporulation delay typical of ECM mutants with significant differences compared to the WT colonies 48 and 72 h. Average values of three biological replicates are shown. Error bars represent the SEM. Statistical significance was assessed by one-way ANOVA with the Dunnet test (*** p<0.001).

**Fig. Sup. 6**

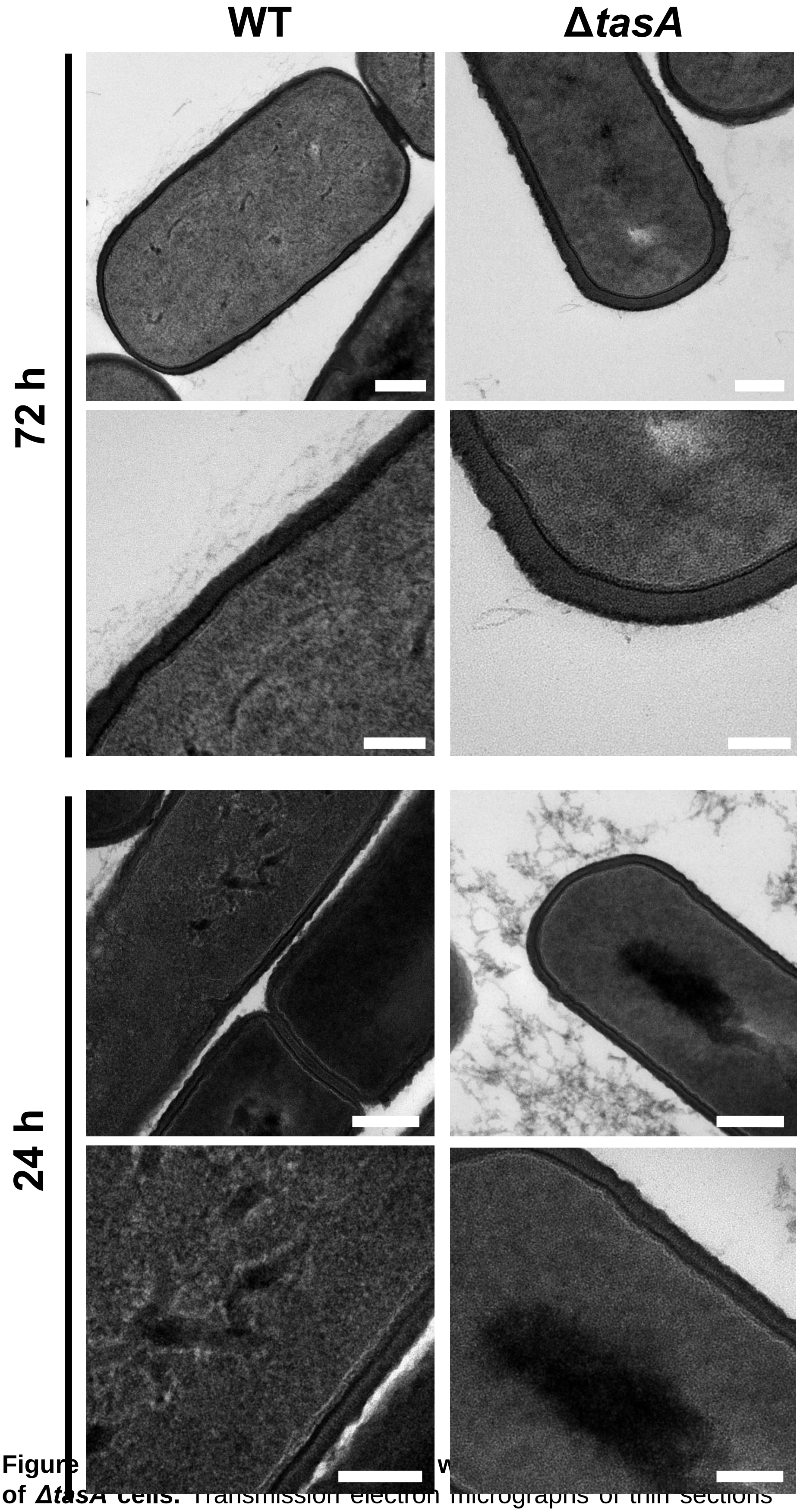


**Figure** [illegible] **of Δ*tasA* cells.** Transmission electron micrographs of thin sections of embedded WT and Δ*tasA* cells at 24 and 72 h post-inoculation. The details of the cell surfaces show no differences in shape, integrity or thickness. Scale bars = 200 nm (top images) or 100 nm (bottom images).

**Fig. Sup. 7**

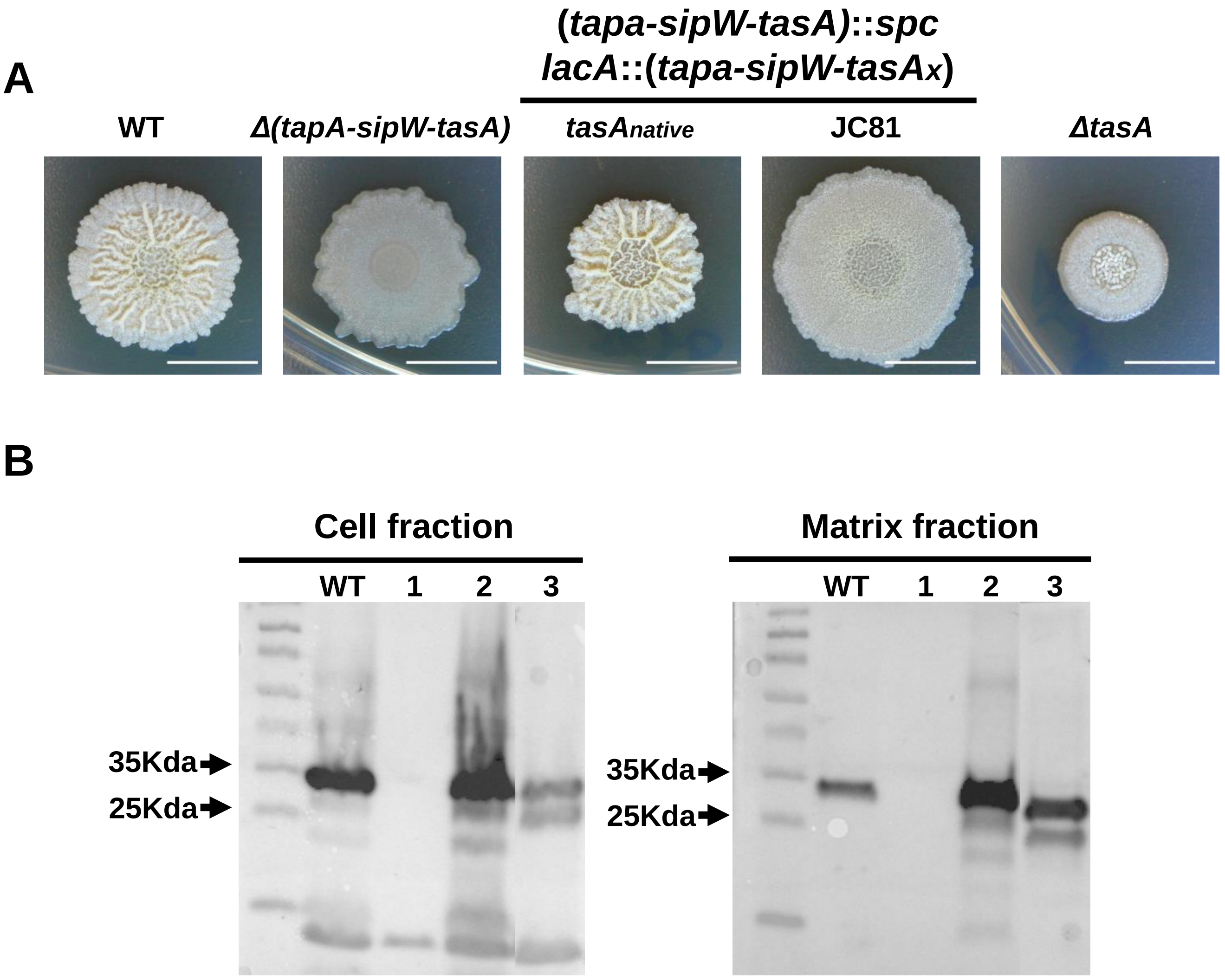


**Figure S7. Phenotypes of the JC81 strain, which carries an allelic variant of *tasA*.** A) A *Δ*(*tapA-sipW-tasA*) strain was used as the genetic background for introducing the entire operon (in *lacA*, a neutral locus) with different versions of *tasA* in which some amino acids were substituted with alanine via site-directed mutagenesis. Colony morphologies of the different *B. subtilis* mutants on MSgg agar plates. From left to right: WT, *Δ*(*tapA-sipW-tasA*), *Δ*(*tapA-sipW-tasA*)::(*tapA-sipW-tasA*$_{\text{native}}$) (containing a WT version of the gene), *Δ*(*tapA-sipW-tasA*)::(*tapA-sipW-tasA*$_{\text{variant}}$) (strain JC81) and *ΔtasA*. B) A western blot of the different biofilm fractions with anti-TasA antibody: cells (left) and ECM (right). Lanes: 1 = WT, 2 = *ΔtasA*, 3 = *tasA*$_{\text{native}}$, and 4 = JC81 (*tasA*$_{\text{variant}}$). Scale bars = 1 cm .

**Fig. Sup. 8**

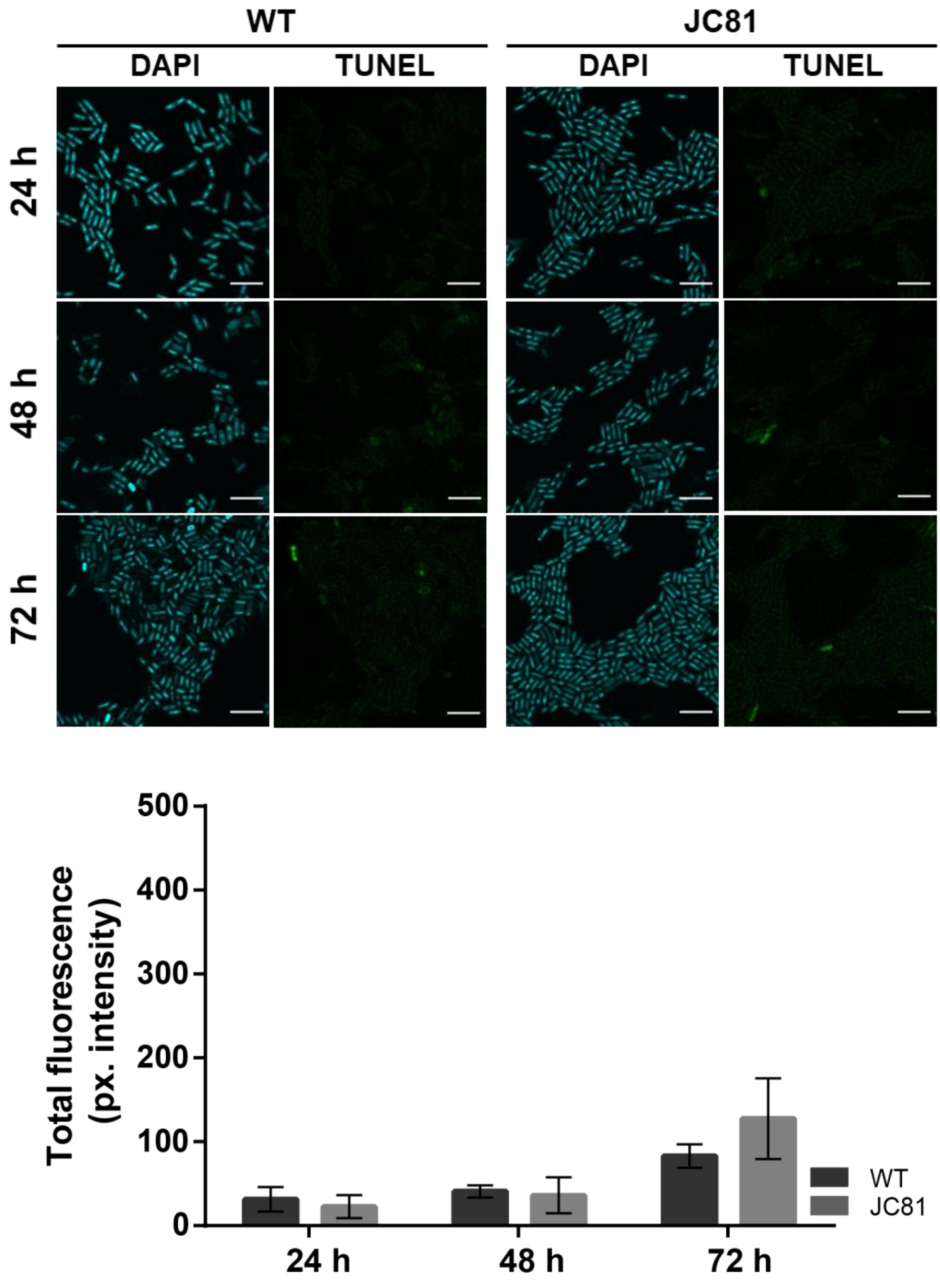


**Figure S8. A TUNEL assay with JC81 showed no difference in the DNA damage level compared with that in the WT strain.** A) TUNEL assay (right) and CLSM analysis revealed no difference in the levels of DNA damage between JC81 colonies and WT colonies. Scale bars = 5 µm. B) Quantification of the TUNEL signals in WT and JC81 colonies showed similar values. Average values of three biological replicates are shown. For each experiment, at least three fields-of-view were measured. Error bars indicate the SEM.

**Fig. Sup. 9**

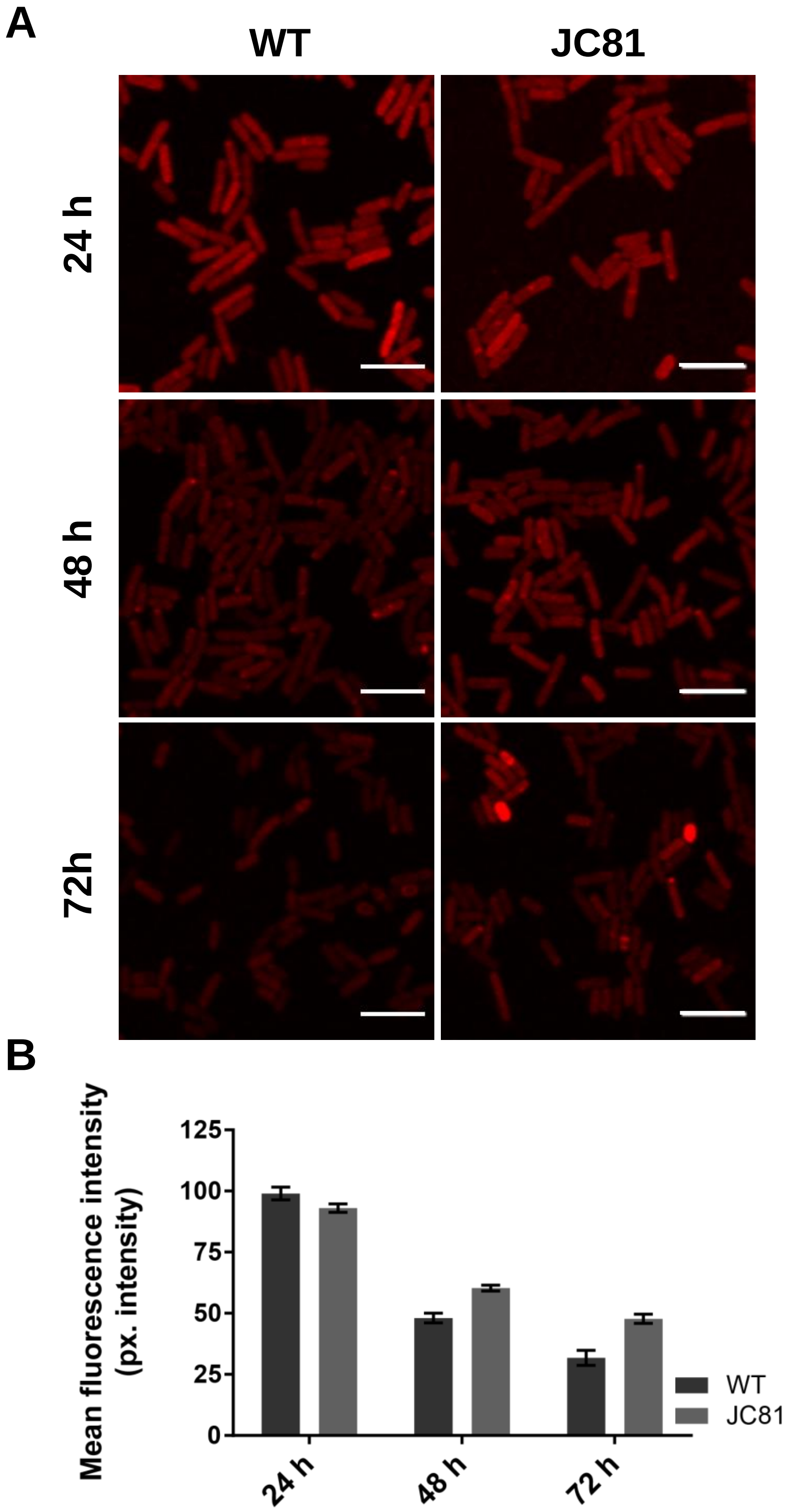


**Figure S9. The membrane potential of JC81 is similar to that of the WT strain.** A) A TMRM assay with WT and JC81 colonies showed no differences in membrane potential. Scale bars = 5 µm. B) Quantification of the TMRM signals at 24, 48 and 72 h post-inoculation showed no differences. Average values of three biological replicates are shown. For each experiment, at least three fields-of-view were measured. Error bars indicate the SEM.

**Fig. Sup. 10**

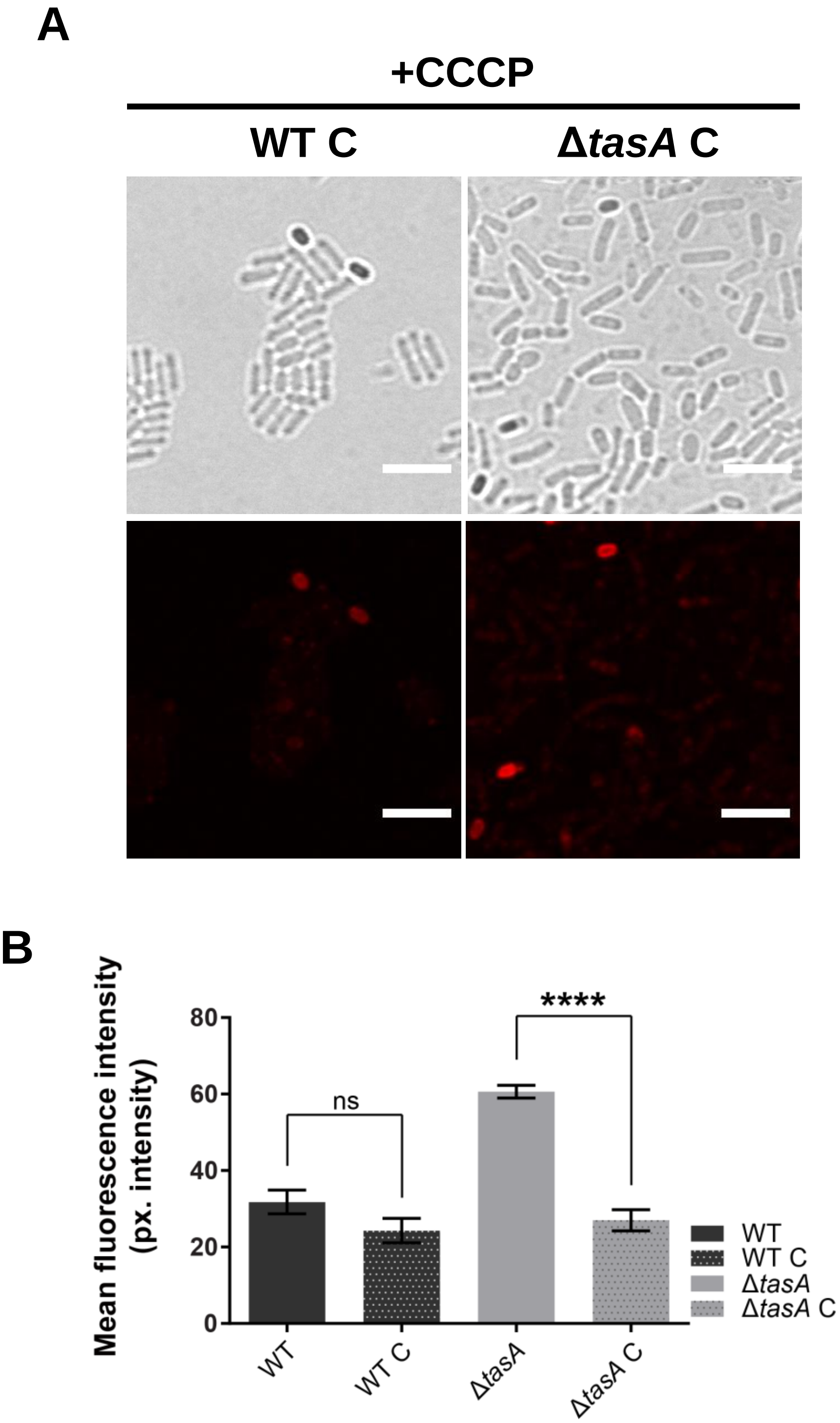


**Figure S10. Treatment of WT and *ΔtasA* cells with a known protonophore caused a decrease in membrane potential.** A) As a control, CLSM analysis of 72-hour-old WT and *ΔtasA* colonies treated with CCCP, a known ionophore that causes proton gradient disruption and depolarization, was performed. The WT cells showed a natural decay in membrane potential over time; therefore, at 72 h, CCCP had very little effect (WT C). However, in the *ΔtasA* cells, which showed hyperpolarization at 72 h, CCCP had a major effect on the membrane potential (*ΔtasA* C), as it could depolarize cells to a level comparable to that in WT C. Scale bars = 5 µm. B) Quantification of the TMRM signals in WT C and *ΔtasA* C showed statistically significant differences in *ΔtasA* C at 72 h compared to that of the untreated sample. Average values of three biological replicates are shown. For each experiment, at least three fields-of-view were measured. Error bars indicate the SEM. Statistical significance was assessed via t-test (*** $p<0.001$).

Fig. Sup. 11

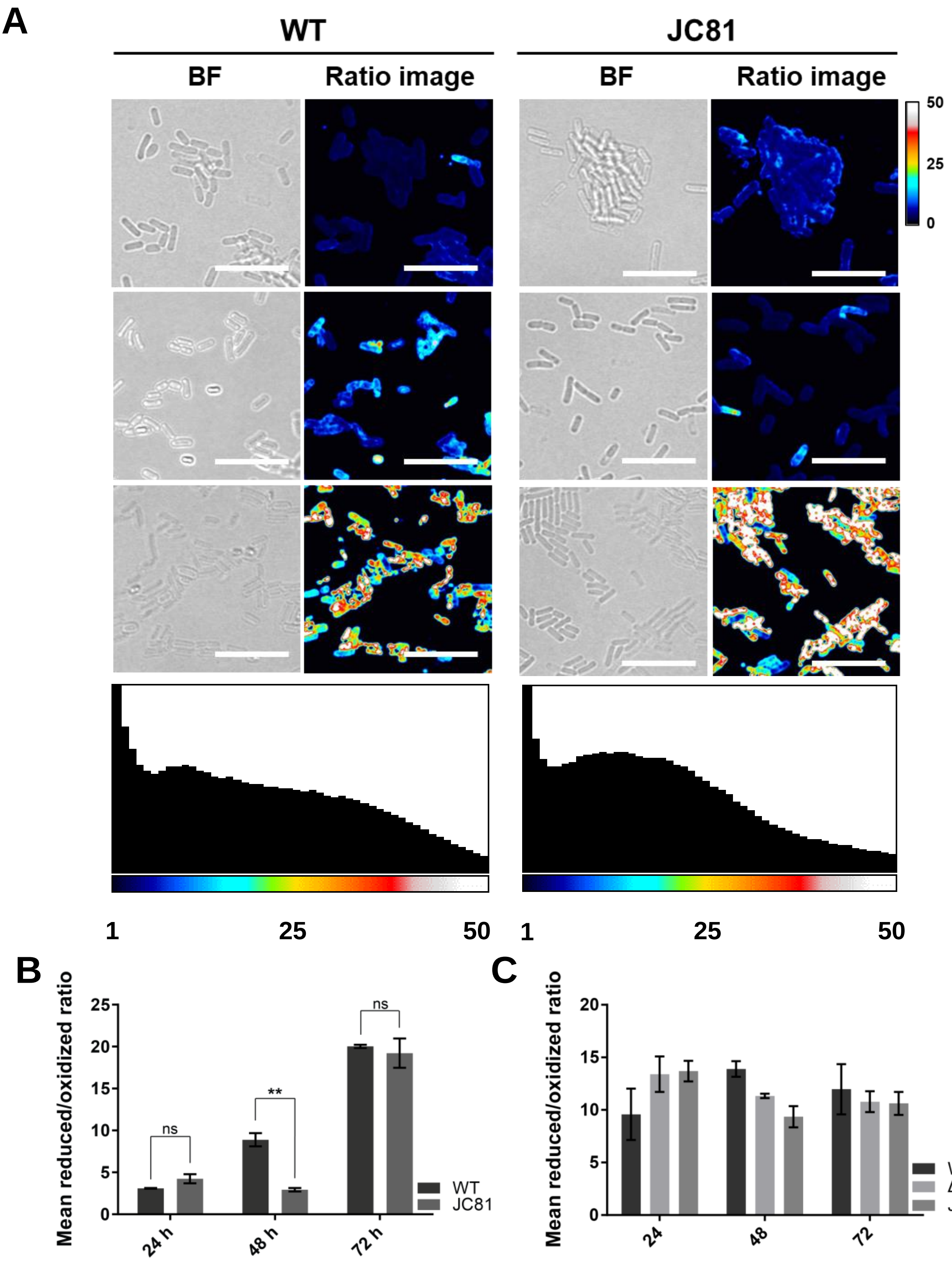


**Figure S11. No differences were found in the susceptibility of the JC81 and WT strains to lipid peroxidation.** A) Assessment of lipid peroxidation in WT and JC81 colonies upon treatment with 5mM CuHpx and analysis by CLSM (top). A calibration bar (from 0 to 50) is located on the right. Confocal microscopy images show that CuHpx treatment was ineffective in both the WT and JC81 strains at 72 h. A histogram of the images shows very little difference in the distribution of pixels based on intensity (bottom) between WT and JC81 at 72 h. Scale bars = 10 μm B) Quantification of the lipid peroxidation from the ratio images shows how the WT and JC81 cells follow the same progression over time after CuHpx treatment, showing increasing values of the reduced/oxidized ratio at the different time points. Average values of three biological replicates are shown. For each experiment, at least three fields-of-view were measured. Error bars indicate the SEM.

**Fig. Sup. 12**

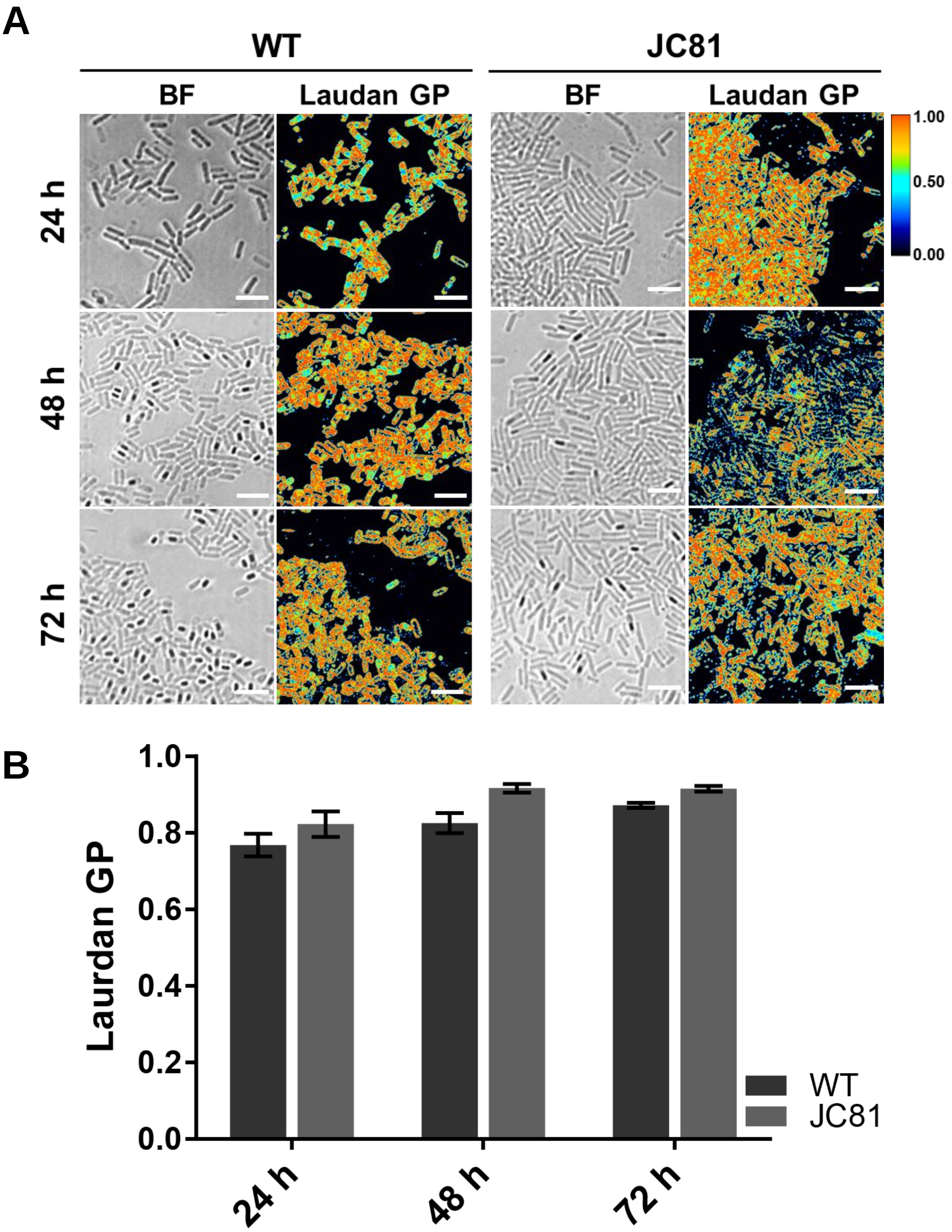


**Figure S12. WT and JC81 cells show similar Laurdan GP values.** A) Laurdan GP analyzed by fluorescence microscopy. A calibration bar (from 0 to 1) is located on the right. The Laurdan GP images show no differences in the membrane fluidities of the WT and JC81 cells at 48 and 72 h. Scale bars = 5 µm B) Quantification of the Laurdan GP showed no difference in the Laurdan GP values between the WT and JC81 cells. Average values of three biological replicates are shown. For each experiment, at least three fields-of-view were measured. Error bars indicate the SEM.

Fig. Sup. 13

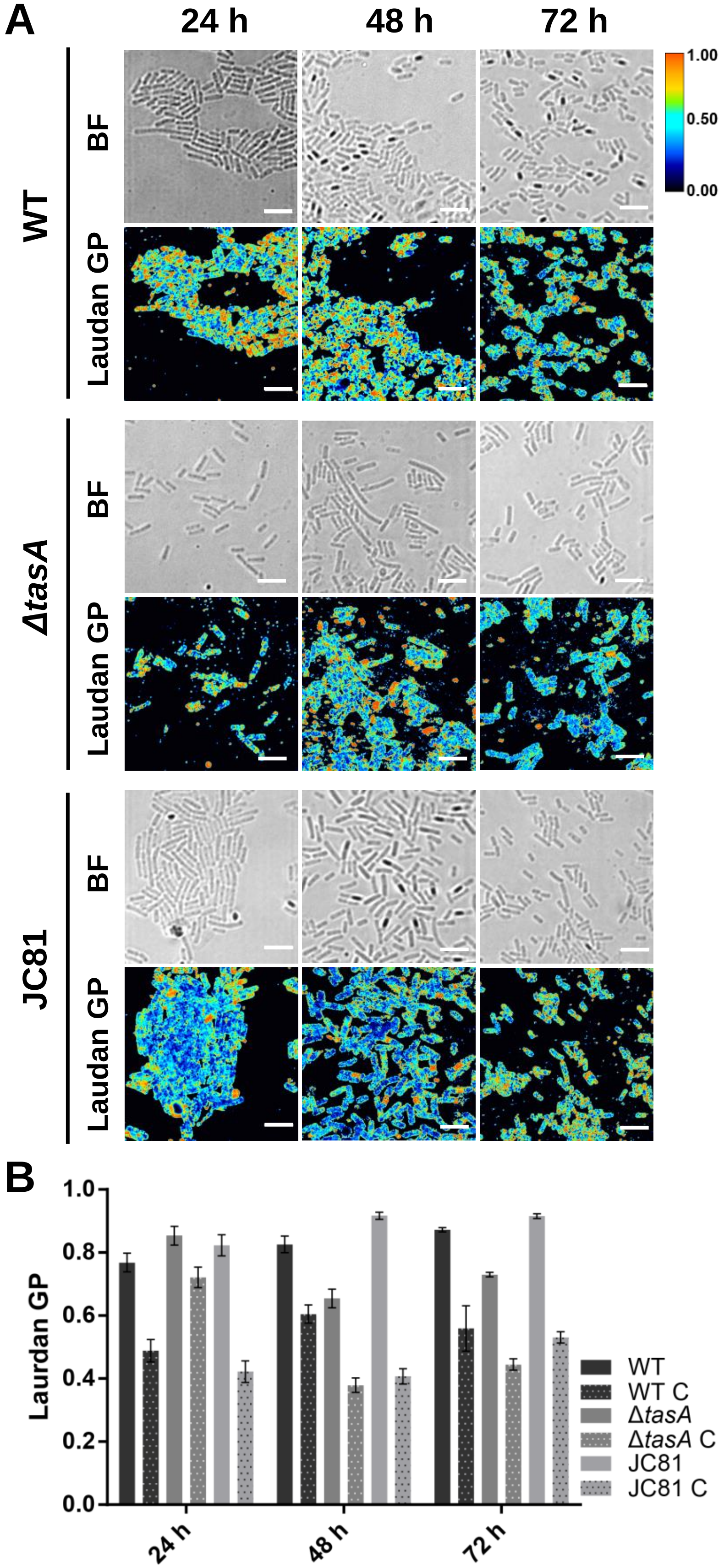


**Figure S13. Membrane fluidity is increased upon treatment of the cells with benzyl alcohol.** A) The Laurdan GP values of WT, *ΔtasA* and JC81 cells 24, 48 and 72 h treated with 2% benzyl alcohol, a known membrane fluidifier, were used as controls. The values were determined using fluorescence microscopy. The images show decreased Laurdan GP values (i.e., increased membrane fluidity) at all time points in all of the strains upon treatment with benzyl alcohol compared with the values of the untreated samples (Fig. 8 and Fig. S13). A calibration bar (from 0 to 1) is located on the right. Scale bars = 5 µm. B) Quantification of the Laurdan GP values in benzyl alcohol-treated samples. Average values of three biological replicates are shown. For each experiment, at least three fields were measured. Error bars indicate the SEM.
